# Morphological, cellular and molecular characterization of posterior regeneration in the marine annelid *Platynereis dumerilii*

**DOI:** 10.1101/352211

**Authors:** Anabelle Planques, Julien Malem, Julio Parapar, Michel Vervoort, Eve Gazave

**Author notes:** = co-senior and corresponding authors (Michel Vervoort, Eve Gazave).

## Abstract

Regeneration, the ability to restore body parts after an injury or an amputation, is a widespread but highly variable and complex phenomenon in animals. While having fascinating scientists for centuries, fundamental questions about the cellular basis of animal regeneration as well as its evolutionary history remain largely unanswered. We study regeneration of the marine annelid *Platynereis dumerilii*, an emerging comparative developmental biology model, which, like many other annelids, displays important regenerative abilities. If the posterior part of the body is amputated, *P. dumerilii* worms are able to regenerate the posteriormost differentiated part of the body and stem cell-rich growth zone that allows to make new segments which replace the amputated ones. We show that posterior regeneration is a rapid process that follows a well reproducible paths and timeline, going through specific stages that we thoroughly defined. Wound healing is achieved by one day post-amputation and a regeneration blastema forms one day later. At this time point, some tissue specification already occurs, and a functional posterior growth zone is re-established as early as three days after amputation. Regeneration is only influenced in a minor manner by worm size and position of the amputation site along the antero-posterior axis of the worm and regenerative abilities persist upon repeated amputations without important alterations of the process. We also show that intense cell proliferation occurs during regeneration and that cell divisions are strictly required for regeneration to normally proceed. Finally, through several 5-ethynyl-2’-deoxyuridine (EdU) pulse and chase experiments, we provide evidence in favor of a local origin of the blastema, whose constituting cells mostly derive from the segment immediately abutting the amputation plane. The detailed characterization of *P. dumerilii* posterior body regeneration presented in this article provides the foundation for future mechanistic and comparative studies of regeneration in this species.

## INTRODUCTION

Regeneration, the replacement of lost body parts, restoring mass and function (Poss, 2010), is a widespread phenomenon in metazoans (Bely and Nyberg, 2009). Regeneration occurs during the life of most or all animals to replace cells that have been lost in day-to-day minor damages, in particular in tissues such as the epidermis and the epithelial lining of the gut (‘homeostatic regeneration’; Poss, 2010). Regeneration can also occur after trauma, such as amputations or ablations (‘injury-induced regeneration’; Poss, 2010). The extant of what can be regenerated after an injury varies a lot among animals: it could be only some cells (or even cell parts such as neuron axons), some tissues (such as the epidermis), some organs (such as the liver), some complex body structures (such as appendages) or even most or all of the body from a piece of tissue (Bely and Nyberg, 2009; Grillo et al., 2016). The ability to regenerate complex body structures, while not found in the most commonly studied animal model such as mammals, *Drosophila melanogaster* and *Caenorhabditis elegans*, is nevertheless found in species that belong to all the main branches of the metazoan tree, raising the possibility that this ability is an ancestral feature of animals and might therefore rely on homologous mechanisms and genetic networks (Bely and Nyberg, 2009).

Two main modes of regeneration have been generally described: epimorphic regeneration that requires active cell proliferation and morphallactic regeneration during which the restoration of the missing body part is solely due to the remodeling of pre-existing cells tissues (Morgan, 1901). In many cases, for example whole-body regeneration in flatworms or limb regeneration in arthropods and vertebrates, epimorphic regeneration involves the formation of a regeneration-specific structure, the blastema, made of a superficial layer of epithelial cells that unsheathes an inner mass of mesenchyme-like cells, and which gives rise to the regenerated structures (Sanchez Alvarado and Tsonis, 2006; Poss, 2010; Tanaka, 2016). A key question in the regeneration field is to determine whether blastemal cells derive from pre-existing (resident) stem cells present in the body before the amputation and ‘activated’ by this injury, or are produced by dedifferentiation of differentiated cells at or close to the amputation site, or a combination of both possibilities (Tanaka and Reddien, 2011). Another crucial question is whether blastemal cells (or at least some of them) have self-renewal capabilities (and could therefore be considered as stem cells) and are pluripotent or multipotent (tissue-restricted) stem/progenitor cells. Planarian regeneration, for example, involves neoblasts (slowly-dividing resident stem cells) which are found throughout the body of the uninjured animals (Sanchez Alvarado and Tsonis, 2006). Neoblasts migrate to the amputation site and produce the regeneration blastema through intense cell divisions. Among the several different categories of neoblasts present in the body of this flatworm, only a specific type, clonogenic neoblasts, were shown to be pluripotent (Reddien, 2013; Wagner et al., 2011). Dependence of regeneration on resident pluripotent stem cells has also been suggested in other animals with whole-body regeneration abilities, such as cnidarians, sponges and colonial ascidians (reviewed in Knapp and Tanaka, 2012; Tanaka and Reddien, 2011). During vertebrate limb regeneration, in contrast, the blastema seems to only contain tissue-restricted progenitor/stem cells, mainly produced by dedifferentiation processes at or near the amputation site, although resident tissue-restricted stem cells (for example satellite cells) may also be involved (Tanaka, 2016).

Annelids (segmented worms) are among the animals that show the most important regenerative abilities (Bely, 2014; Özpolat and Bely, 2016). Many annelids are able to regenerate, after an amputation, the posterior part of their body, their anterior part (including the head), or both. There is a long history of experimental and descriptive studies of regeneration in many annelid species and more recently cellular and molecular aspects of this process have been studied in a few species, including *Alitta virens*, *Pristina leydyi*, *Capitella teleta*, *Eisenia fetida*, and *Enchytraeus japonensis* (for review; Bely, 2014; Özpolat and Bely, 2016). Most of these species belong to one of the two main clades of the phylum Annelida, the Sedentaria (Struck et al., 2011). In these species, indirect evidence for an involvement of neoblast-like resident stem cells during regeneration, such as the presence in non-injured animals of cells that express homologues of genes known to be expressed in flatworm neoblasts and the observation of long distance migration of cells towards the amputation site, have been obtained (*e.g.,* Bely, 2014; de Jong and Seaver, 2017; Myohara, 2012; Özpolat and Bely, 2016; Zattara et al., 2016). In the other main annelid group, the Errantia, the presence of neoblast-like cells is much more elusive and there are in contrast experimental evidence indicating that regenerated structures only originate from cells located close to the amputation site, probably through dedifferentiation events (*e.g.*, Boilly, 1965a,b; Boilly, 1968a,b; Boilly, 1969a,b,c).

In this study, we investigated regeneration of the marine annelid worm *Platynereis dumerilii* (Audouin and Milne Edwards, 1833), which belongs to the Errantia clade, and which has proven to be very useful for large-scale evolutionary developmental comparisons (Raible and Tessmar-Raible, 2014; Williams and Jékely, 2016). *P. dumerilii* displays a complex life cycle (Fischer et al., 2010): embryonic and larval development lead in three days to the formation of a three-segmented worms that will subsequently grow during most of their life, by sequentially adding new segments at their body posterior end, just before the pygidium. This process, known as posterior growth, relies on the presence of a subterminal ‘growth zone’ that contains stem/progenitor cells expressing a complex molecular signature, the Germline Multipotency Program (GMP; Juliano et al., 2010), also displayed by pluripotent/multipotent somatic stem cells and primordial germ cells in other animals (Gazave et al., 2013). *P. dumerilii* worms also have important regeneration abilities: after amputation of the posterior part of their body, which leads to the removal the pygidium (final non-metameric part of the body), the growth zone and several segments, the worms are able to regenerate the pygidium and the growth zone which will in turn produce new segments (Gazave et al., 2013). In this article, we performed a detailed characterization of this regeneration process (that we named posterior regeneration) at the morphological, cellular and molecular levels. We found that *P. dumerilii* posterior regeneration is a rapid process that passes through well-defined and reproducible stages. Regenerative abilities are conserved after multiple repeated amputations. Posterior regeneration involves the formation of a blastema-like structure and requires cell proliferation, therefore being of the epimorphic type. EdU (5-ethynyl-2’-deoxyuridine, a thymidine analog) pulse and chase experiments do not support the hypothesis that *P. dumerilii* posterior regeneration may rely on the long-distance migration of stem cells and strongly suggest in contrast that the regenerating region is mostly produced by cells of the differentiated segment abutting the amputation plane.

## MATERIALS AND METHODS

### Breeding culture and worm collection

*P. dumerilii* worms were obtained from a breeding culture established in the Institut Jacques Monod, according to the protocol of Dorresteijn et al. (1993). In most experiments, 3-4 months’ worms with 30-40 segments were used. After anesthesia (with a solution of MgCl2 7.5 % and sea water, 1/1), sharp amputations of the posteriormost part of the worms were done using a microknife (Sharpoint™) between two segments (perpendicularly to the body axis), in order to remove the 5 to 6 posteriormost segments and the pygidium. After amputation, worms were let to recover in fresh sea water and fed normally three times per week. For serial amputations, worms were amputated the 1^st^ time as explained above and the following amputations were performed to remove the regenerated region plus the previous non-amputated segment. At defined time points after amputation, worms were fixed in 4% PFA in 100mM PBS tween 0.1% (PTW) for 2 hours at RT, washed with PTW and stored at −20°C in 100% MeOH for subsequent experiments. For phalloidin staining (see below), worms were similarly fixed in PFA and rinsed in PTW but were then directly stored at 4°C for up to 4 days before labeling, without MeOH dehydration.

### Micro-computed X-ray tomography (microCT) and scanning electron microscopy (SEM)

For microCT imaging, specimens were gradually dehydrated in ascending ethanol series up to 96% ethanol and then dehydrated for two hours with hexamethyldisilazane (HMDS) and left to dry overnight before scanning, following Alba-Tercedor & Sánchez-Tocino (2011). Scanning was carried out with a microtomograph Skyscan 1172 (Bruker, Belgium) using the following parameters: voltage 55 kv, current 165 uA, and pixel size between 0.54 and 1.28 um depending on the sample size. No filter was used, and samples were rotated 360° for obtaining as much detail as possible. The X-ray projection images obtained during scannings were reconstructed with the software NRecon (Bruker, Belgium). The sections obtained were processed with the programmes CTAn and DataViewer (Bruker, Belgium). 3D representations were obtained with the programme CTVox (Bruker, Belgium). For Scanning Electron Microscopy (SEM) experiments, specimens were dehydrated *via* a graded ethanol series, prepared by critical-point drying using CO_2_, mounted on aluminium stubs, covered with gold in a BALTEC SCD 004 evaporator, and examined and photographed under a JEOL JSM-6400 SEM.

### *Cloning of* Pdum-piwiB *and phylogenetic analysis of piwi genes*

A previously unidentified *P. dumerilii piwi* gene (subsequently named *Pdum-piwiB*) was found by sequence similarity searches against the *P. dumerilii* transcriptomic database PdumBase (pdumbase.gdcb.iastate.edu; Chou et al., 2016). The gene was cloned using standard protocols as described in Gazave et al. (2013). Phylogenetic tree reconstruction was performed using the dataset of *piwi/argonaute* genes used by Kerner et al. (2011) to study the evolution of this large gene family in metazoans, and methods described in Gazave et al. (2013).

### *Whole-mount* in situ *hybridizations (WMISH), immunolabelings, EdU cell proliferation assay, and imaging*

NBT/BCIP WMISH and immunolabelings were performed as previously described (Tessmar-Raible et al., 2005; Demilly et al., 2013; Gazave et al., 2017). For all labelings, after rehydration, samples were treated with 40 μg/ml proteinase K PTW for 10min, 2mg/ml glycine PTW for 1min, 4% PFA PTW for 20min and washed in PTW prior to hybridization or labeling. Neurites, as well as cells undergoing mitosis and proliferating cell nuclear antigen labelings were done as previously described (Demilly et al., 2013), using the mouse anti-acetylated tubulin (Sigma T7451, 1:500), the rabbit anti-phospho histone H3 (Milipore 06-570, 1:100) and the PCNA PC10 (sc-56, Santa Cruz Biotechnology inc., 1:500) antibodies, respectively, and fluorescent secondary anti-mouse IgG Alexa Fluor 488 or 555 conjugate (Invitrogen, 1:500). For phalloidin labeling, digestion and post-fixation steps were done similarly, on non-rehydrated samples, that were subsequently incubated in phalloidin-Alexa 555 (Molecular Probes, 1:100) overnight with shaking at 4°C.

Proliferating cells were labeled by incubating worms with the thymidine analog 5-ethynyl-2′-deoxyuridine (EdU), which was subsequently fluorescently labeled with the Click-It EdU Imaging Kit (488 or 555, Molecular Probes) as described in Demilly et al. (2013). Worms were incubated for 1h or 5h in sea water with 5 μM EdU. Various pulse and chase experiments were performed as described in the result section.

Following fluorescent labeling procedures, samples were counterstained with Hoechst (or DAPI) at 1:1000 dilution and stored and mounted in 87% glycerol containing 2.5 mg/mL of anti-photobleaching reagent DABCO (Sigma, St. Louis, MO, USA) before confocal imaging.

Bright field images were taken on a Leica microscope for visible NBT/BCIP WMISH. Confocal images were taken with a Zeiss LSM 710 confocal microscope. Adjustment of brightness and contrast, Z projections were performed using ImageJ and Adobe Photoshop. The figure panels were compiled using Adobe Illustrator.

### Cell proliferation inhibitor treatments

Cell proliferation during regeneration was blocked using two well characterized inhibitors, hydroxyurea (HU, Sigma H8627) and nocodazole (Sigma M1404). Nocodazole treatments, even at low concentration (0.01μM, 0.1μM and 1μM), while blocking regeneration, induced high worm autotomy and lethality (not shown) and therefore further experiments were done only with HU. HU was dissolved in sea water (10, 20mM or 50mM) and the HU solution was changed every 24 h to maintain its activity for the duration of the experiment. Worms were incubated in 2ml of HU solution in 12-wells plate for the desired time period (see Results for details).

### Scoring and statistical analyses

For morphological and HU treatment experiments, worms were observed under the dissecting microscope at different time points after amputation and ‘scored’ according to the regeneration stage that has been reached (stages 1 to 5; see Results and Fig. 3 for the definition of stages) or the number of morphologically well visible segments that have been produced (X s., X being the number of visible segments). Some worms showed a morphology that was intermediate between that of two successive stages and were therefore scored as 1.5, 2.5, 3.5, and 4.5.

Graphic representations of morphological experiments and statistical analyses were performed using the Prism 7 software (GraphPad). For groups multiple comparison (worm sizes and position of amputation experiments; Fig. 4), 2-way ANOVA on repeated measures using Tukey correction were used. For the study of worms’ regeneration with daily scoring after four serial amputations (Fig. 5), 2-way ANOVA on 2 factors repeated measures using Dunnett correction was performed. Two worms died during the experiment and were thus removed from analysis as well as well one worm whose score was missing for one day. For HU treatments (Fig. 10), 2-way ANOVA on repeated measures using Dunnett correction was performed. Comparison with similar p-values were grouped for graphic representation.

## RESULTS

### *Definition of* P. dumerilii *posterior regeneration stages*

We tested the regeneration abilities of juvenile worms that are in the post-larval growth phase of their life cycle during which they grow by adding one by one new segments in the posterior part of their body (posterior elongation). This phase starts at the end of embryonic and larval development with small worms displaying a head, three segments with a pair of lateral appendages (parapodia), and a terminal body non-segmental element called the pygidium, and ends when the worms become sexually-mature. In our worm culture conditions, this growth phase spans over a maximum of one year (its duration varies however much from one individual to the other) and, at its end, the worms are made of more than 80 segments. While the number of segments of the worms increases with age, there is no strict correlation between the number of segments, the overall size and the age of the worms.

In a first set of experiments, we removed, from worms long of 30-40 segments (3-4 month-old), the pygidium (which bears the anus and characteristic bilateral outgrowths named anal cirri) and the last five to six posterior segments. We then characterized their regeneration using scanning electron microscopy (SEM; Fig. 1), micro-computed X-ray tomography (microCT; Fig. 2), and bright field microscopy (Fig. 3A). We found that regeneration proceeds through five reproducible stages with a similar timing in most of the worms (Fig. 3B,C). At 1-day post amputation (1dpa), the amputation surface is covered by a wound epithelium but there is no sign of any outgrowth of tissues (stage 1; Figs. 1B and 3A). At 2dpa, a small posterior protuberance (regenerated region) is observed with a depression on its ventral side that probably corresponds to the position of a reformed anus. A loosely organized gut-like structure can be seen in the regenerated region, but no other differentiated internal structures are found (stage 2; Figs. 1C, 2B-B’ and 3A). At 3dpa, the regenerated region has clearly increased in size and very small anal cirri are observed. A well-differentiated gut is present (stage 3; Figs. 1D, 2C-C’ and 3A). The size of the regenerated region continues to increase during the two following days. At 4dpa, a well differentiated pygidium with long anal cirri is present (stage 4; Figs. 1E and 3A). At 5dpa, small lateral indentations separate the pygidium from the more anterior part of the regenerated region in which faint segmental grooves started to be seen. The segmental ganglia of the ventral nerve cord and well defined ventral body wall muscle layer are now visible in the regenerated region (stage 5; Figs. 1F, 2D-D’ and 3A). During the following days, segments with well visible boundaries and developing parapodia can be seen and their number increases rapidly. The number of morphologically well visible segments varies among the worms and at 10dpa ranges from 4 to 11 (Fig. 3B,C).

**Figure 1:**
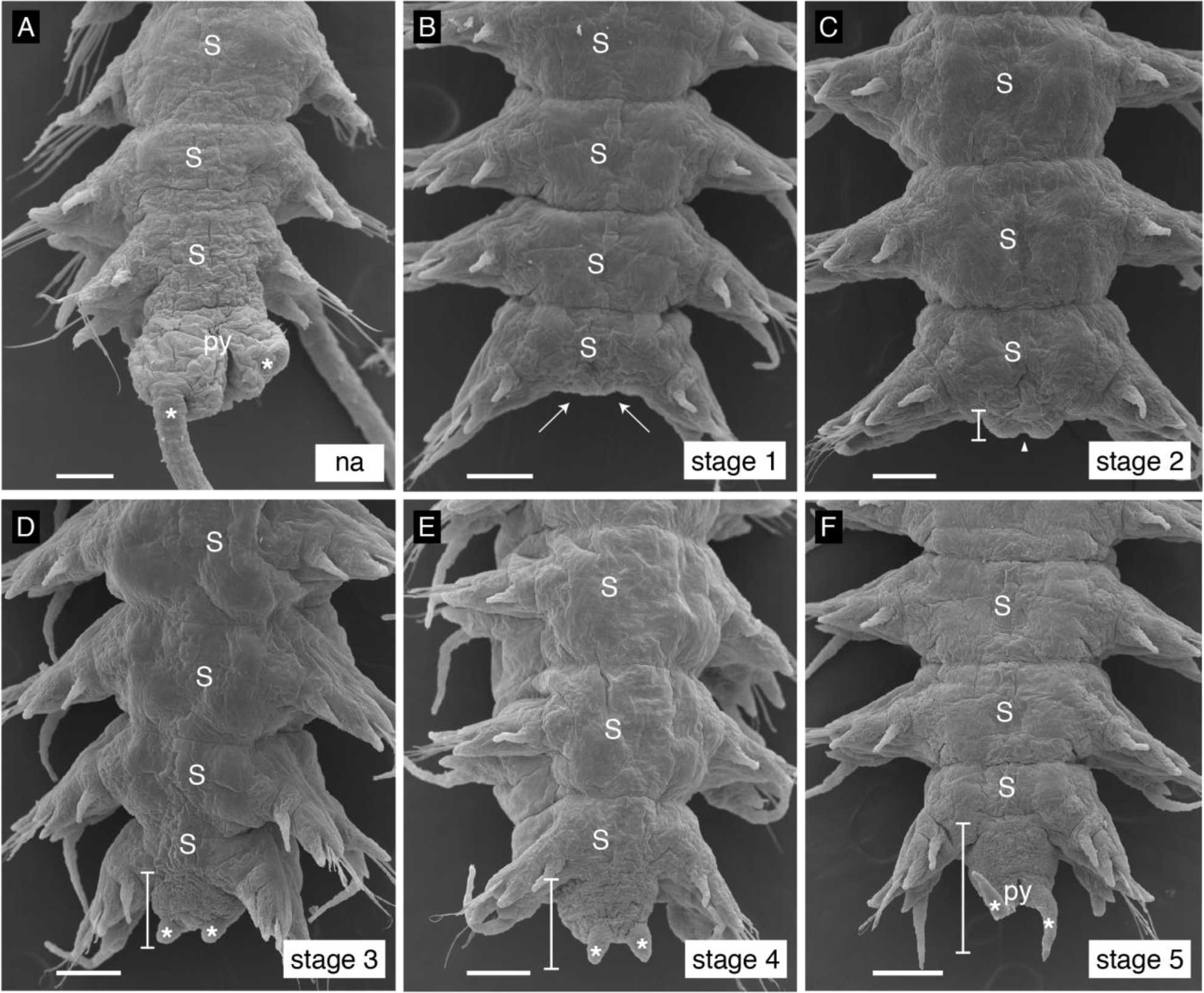
Scanning Electron Microscopy (SEM) micrographs of posterior regeneration stages. (A) Non-amputated (na) condition showing the posterior part of a worm including the last three body segments (S) and the pygidium (py) bearing the anus and anal cirri (asterisks). Left anal cirrus is detached (right side, out of focus). (B) Stage 1 (1-day post amputation, 1dpa): wound healing is achieved (white arrows) but no posterior outgrowth is present. (C) Stage 2 (2dpa): a small regenerated region is visible (white bracket) with a notch in its central part (white arrowhead) that likely corresponds to the reformed anus. (D) Stage 3 (3dpa): regenerated region has increased in size, and two small anal cirri are already visible (asterisks). (E) Stage 4 (4dpa): size of the regenerated region and anal cirri has increased as compared to the previous stage. (F) Stage 5 (5dpa): an indentation separates the differentiating pygidium (bearing longer anal cirri) from the anterior part of the regenerated region. Right ventral cirrus is ventrally bent. All pictures are ventral views. Scale bars= 50 μm. White brackets delineate the regenerated region. S= segment; py= pygidium; asterisks= anal cirri; arrowhead = anus; arrows = wound epithelium.

**Figure 2:**
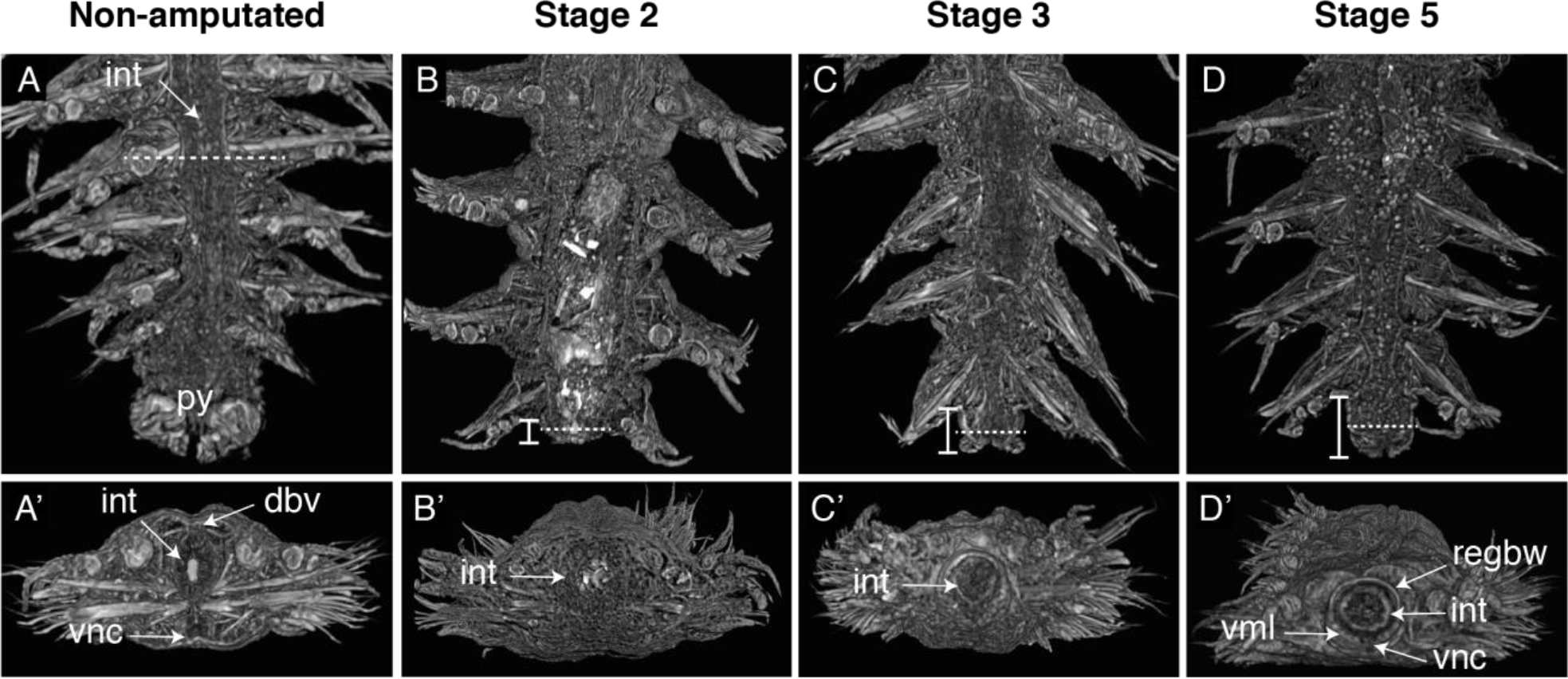
Micro-computed X-ray tomography (microCT) images of posterior regeneration stages. A to D images correspond to mid-coronal sections of the posterior part of a non-amputated worm (A) and at different stages of regeneration (B-D). The dotted lines correspond to the virtual plan of the transversal sections shown in A’ to D’. Brackets highlight the regenerated region in B to D. int= intestine; py= pygidium; dbv= dorsal blood vessel; regbw= regenerated body wall; vml= ventral muscular layer; vnc= ventral nerve cord. (A-A’) In a non-amputated worm, several internal structures such as the intestine, dorsal and ventral blood vessels and ventral nerve cord can be seen. (B-B’) and (C-C’) At stage 2 and 3, only the intestine is distinguishable in the still small regenerated region. (D-D’) At stage 5, the regenerated region contains well-defined intestine, ganglia of the ventral nerve cord and ventral wall muscle layers.

**Figure 3:**
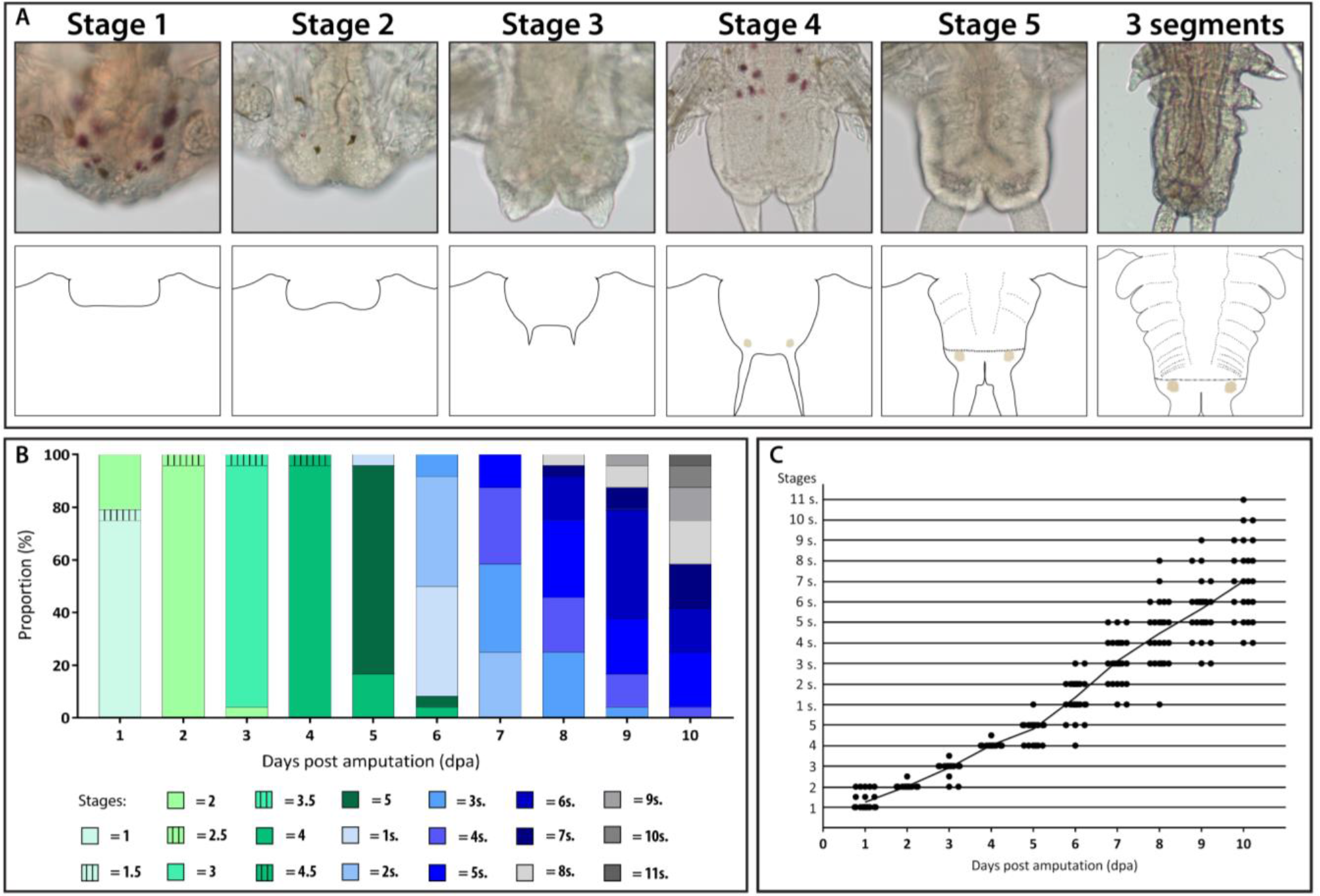
Posterior regeneration proceeds through morphologically well-defined stages with a reproducible timing. (A) Bright-field microscopy images and schematic representations of the five stages of regeneration, as well as an example of a later stage at which three new well-visible segments have been added (and therefore named stage 3 segments; noted 3s. in graphics). (B and C) 27 worms of 3-4 months and with 30 to 40 segments were amputated (removal of the last 5-6 segments and the pygidium) and scored for the reached stage every day for 10 days. (B) Diagram showing the proportion of worms at a specific stage after amputation and follow up for 10 days. Each stage is represented by a color code. Early stages are in green and later stages are in blue and gray. Worms showing a morphology intermediate between that of two successive stages were scored as x.5 (for example 1.5 denote a morphology intermediate between those of stages 1 and 2). Early stages (stage 1 to 5) are in green, later stages at which there are visible segments are in blue and grey and scored according to the number of visible segments (*e.g.*, 1s = 1 visible segment). (C) Graphic representation of the stages reached by the worms every day during 10 days. Each dot corresponds to one individual. It is clear from (B) and (C) that the timing of early stages (stage 1 to 5), corresponding to regeneration *per se*, is very similar in all worms and that these stages correspond to one to five days after amputation. Interindividual variability becomes higher at later stages, corresponding to the phase of segment addition.

### *Parameters influencing* P. dumerilii *posterior regeneration*

We studied parameters that may affect posterior regeneration efficiency and timing. For all these experiments, the worms were ‘scored’ at different time points after amputation by defining the regeneration stage that they have reached (stages 1 to 5) for early time points, or the number of morphologically well-defined segments that have been produced for later time points. We first tested the influence of worm size by comparing regeneration of worms with three different segment number ranges, 10-20 segments (‘small worms’), 30-40 segments (‘intermediate worms’, previously used to define the regeneration stages) and 70-80 segments (‘big worms’; Fig. 4A). Efficient regeneration with the aforementioned reproducible stages was observed in the three categories of worms. While the timing of the process was similar in intermediate and big worms, small worms regenerated and produced segments significantly faster than the other ones (Fig. 4A). Interindividual variability differed in the three worm categories and was the lowest in the 30-40 segment worms (Fig. 4B) that were therefore used for all next experiments.

**Figure 4:**
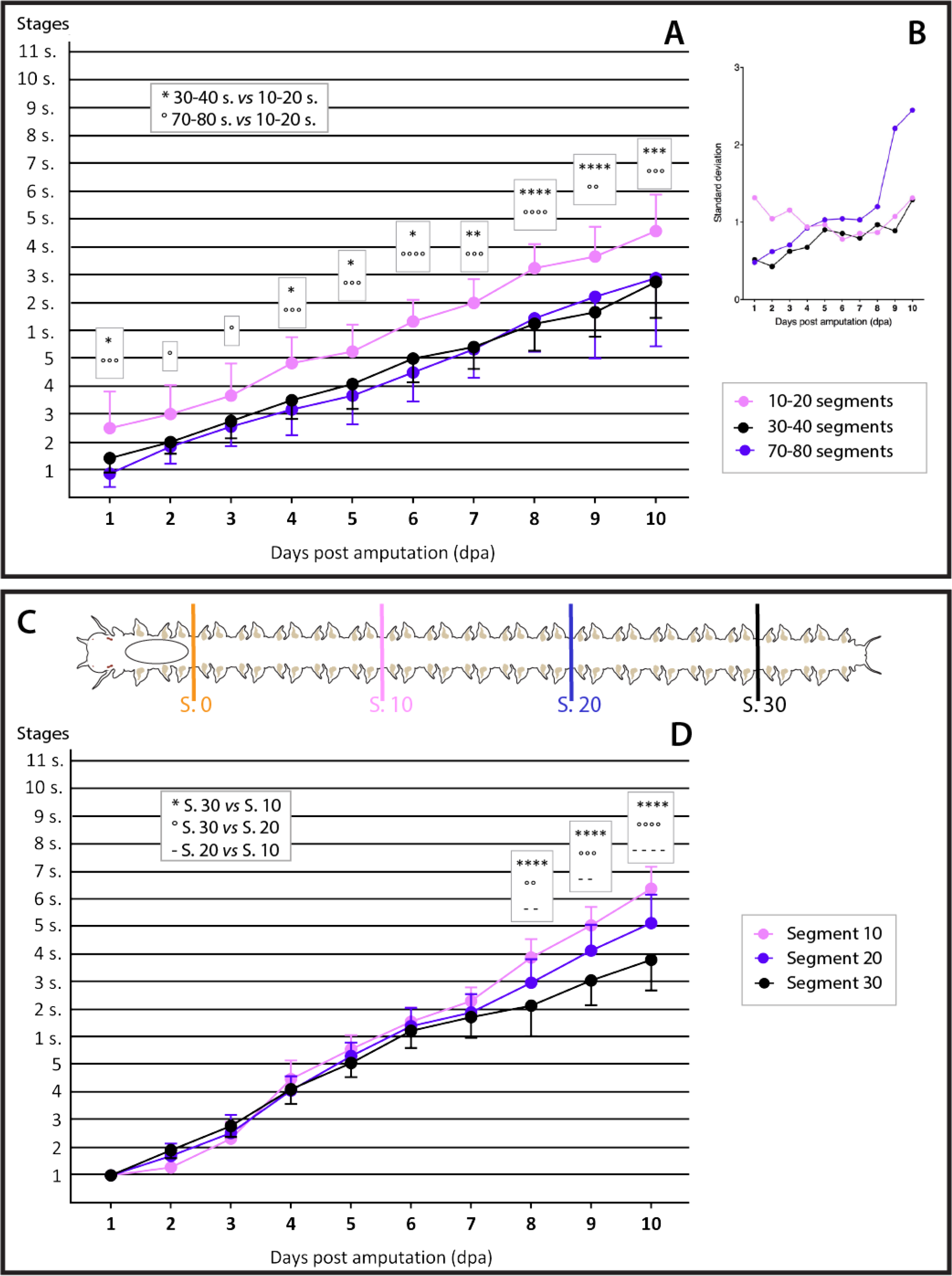
Influence of worm size and position of the amputation plane on posterior regeneration. (A-B) Influence of worm size on posterior regeneration. Three categories of worms were used: worms with 10-20 segments (in pink, n=12), worms with 30-40 segments (in black, n=12) and worms with 70-80 segments (in blue, n=18). (A) Graphic representation of the stages reached by the worms every day during ten days. Whereas complete regeneration and production of segments are observed in all three conditions, worms with 10-20 segments regenerate and produce segments significantly faster than the longer worms. 30-40 segments and 70-80 segments worms do not show significant difference in their timing of regeneration. (B) Graphic representation of the standard deviation (SD, used as a measure of interindividual variability) of the stages reached by the worms during the 10 days following amputation. The worms with 30-40 segments show the lowest variability during the whole process. (C and D) Influence of amputation plane position on posterior regeneration. (C) Schematic representation of the four different positions of amputation along the worm body axis that have been tested. The position 0 corresponds to the location immediately posterior to the pharynx (schematized as an ellipse). Positions 10, 20 and 30 correspond to the number of segments posteriorly to this position 0. In (C) and (D), each amputation position is color-coded as follow: position 0= orange, position 10= pink, position 20= blue and position 30= black. (D) Graphic representation of the stages reached by the worms every day during ten days. Worms amputated at position 0 died in 1 or 2 days without any sign of regeneration and are therefore not represented. Regeneration and segment addition occurred in the three other conditions. There is no significant difference in the timing of the process during the seven first days. From day 8, in contrast, worms amputated more anteriorly produce more segments than those amputated in more posterior positions. Statistics for (A) and (D): 2-way ANOVA on repeated measures using Tukey correction were done. * p<0.05; ** p<0.01; *** p<0.001; **** p<0.0001. Error bar: SD

We next assessed the influence of the position of the amputation site, by comparing four different ones as shown in Fig. 4C (segment 30 corresponds to the amputation site used in the previous experiments). Worms amputated at segment 0 (immediately posterior to the pharynx) died in one or two days without any sign of regeneration and showed an eversion of the pharynx through the amputation site, which could be the cause of the death (not shown). Worms amputated at segment 10, 20 and 30 regenerated in a very similar way (Fig. 4D), indicating that similar regeneration capabilities are found along most of the animal body axis and therefore are not specific of the posterior part of the body. Interestingly the rate of segment addition depends on the position of the amputation and is significantly higher if this position is more anterior (Fig. 4D).

We finally tested how regeneration is impacted by repeated amputations. In a first experiment, we performed four serial amputations with intervals of ten days (at that time the worms have already regenerated the pygidium and produced several segments) and scored the worms every day during the whole experiment (Fig. 5A). Regeneration capability was maintained during these four occurrences of regeneration with only minor modifications of its timing (Fig. 5B). Interindividual variability became however high after the fourth amputation (Fig. 5C). We next put the worms in a more challenging situation: we performed ten serial amputations with intervals of only four days, meaning that new amputations were made before the worms have started again to produce segments, and we scored the worms every four days just before to make the new amputation (Fig. 5D). We found that most worms normally regenerate after all serial amputations (Fig. 5E). There was no clear tendency for either an increase or a decrease of the speed of the process, but however interindividual variability tended to increase (Fig. 5F) and very few worms showed abnormally shaped pygidium and anal cirri after several amputations (not shown).

**Figure 5:**
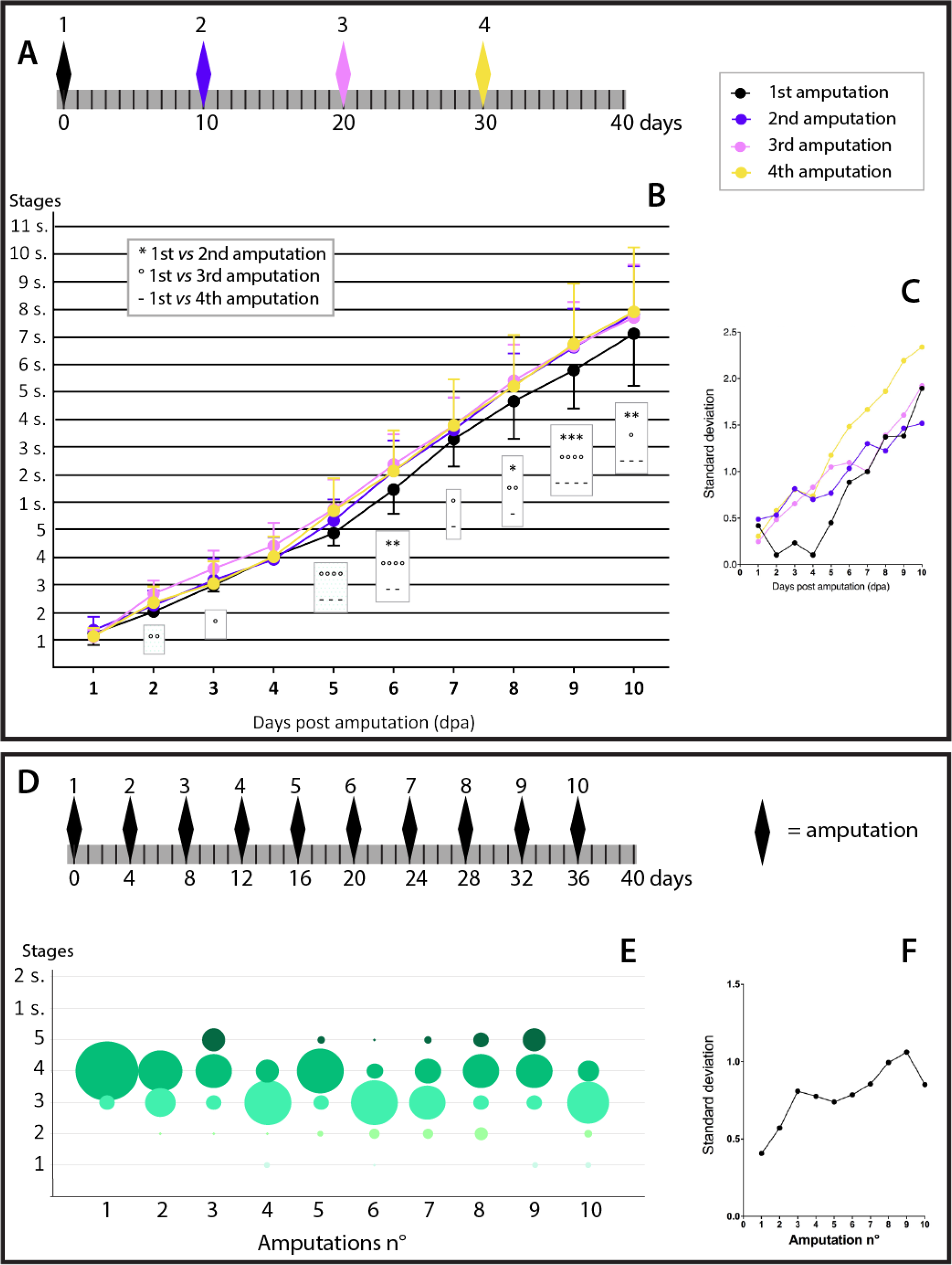
Posterior regeneration abilities are maintained after multiple amputations. (A) Schematic representation of the experiment with four serial amputations performed every ten days. A time line of 40 days is represented in gray with a black bar for each day. Scoring was done every day. Each amputation is represented by a diamond of a specific color as follow: first amputation in black, second amputation in blue, third amputation in pink and fourth amputation in yellow. The same color code is used in (B) and (C). (B) Graphic representation of the stages reached by the worms every day during 10 days after each of the four successive amputations. Efficient regeneration is observed after the four serial amputations and tends to be slightly faster after the second, third and fourth amputations as compared to the first one. At later stage, variability increases. (n=24) (C) Graphic representation of the SD (used as a measure of interindividual variability) of the stages reached by the worms during the 10 days following amputation after each of the four successive amputations. Variability tends to increase after the successive amputations and becomes very important after the fourth one. (D) Schematic representation of the experiment with 10 serial amputations every four days. Each amputation is represented by a black diamond. Scoring were done just before performing a new amputation. (E) Bubble chart representation of the stages reached by the worms four days after each of the 10 amputations. Bubble diameters are proportional to the number of worms at a specific stage. Color-code of the bubble is that of Figure 3. Regeneration abilities are maintained after 10 serial amputations and there is no clear tendency for either an increase or a decrease of the speed of the process. (n=30) (F) Graphic representation of the SD of the stages reached by the worms four days after each of the 10 successive amputations. Variability tends to increase during the successive amputations. Statistics for (B): 2-way ANOVA on 2 factors repeated measures using Dunnett correction was performed. * p<0.05; ** p<0.01; *** p<0.001; **** p<0.0001. Error bar: SD

We therefore conclude that *P. dumerilii* posterior regeneration passes through reproducible and well-defined stages and is only influenced in a minor manner by worm size and position of the amputation site along the antero-posterior axis of the worm. Regenerative abilities persist upon repeated amputations without important alterations of the process, even after ten successive and close in time amputations.

### *Molecular and cellular characterization of* P. dumerilii *posterior regeneration*

To further characterize *P. dumerilii* posterior regeneration, we performed whole-mount *in situ* hybridizations (WMISH) for several genes previously shown to be involved in segment, organ or tissue patterning and differentiation in *P. dumerilii* (Table S1). Expression patterns of all these genes were previously established during posterior elongation in about one-year-old worms 10 to 12 days after posterior amputation when many segments have already been produced (Gazave et al., 2013). We found very similar expressions in 30-40 segment (3-4-month-old) worms five days after posterior amputation (5dpa, stage 5; Fig. S1). Figure 6 shows representative images (mainly ventral views) of the expression of all the genes at the earlier stages (stage 1 to 4). Some additional images (mainly dorsal views) highlighting other interesting aspects of the expression of some of the genes at some stages are shown in figure S2.

**Figure 6:**
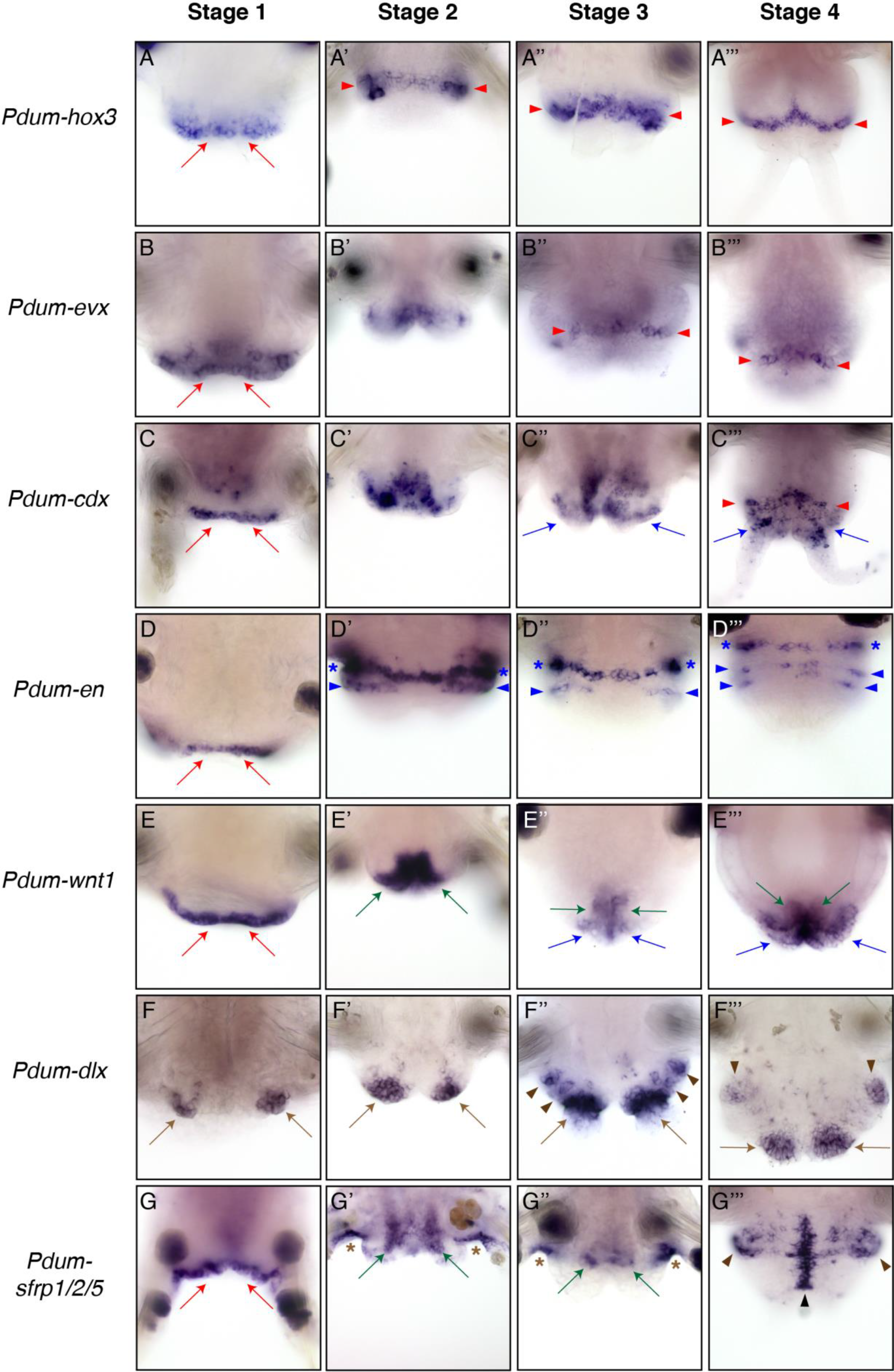
Expression of genes involved in segment, organ or tissue patterning and differentiation during posterior regeneration. Whole-mount *in situ* hybridizations (WMISH) for the genes whose name is indicated are shown for four posterior regeneration stages (stage 1 to 4). Stage 5 WMISH images are shown in Fig. S1. All panels are ventral views (anterior is up). Red arrows = wound epithelium; brown arrows = anal cirri; green arrows = intestine; blue arrows = pygidium; black arrows = neurectoderm or ventral nerve cord; pink arrows = pygidial muscles; red arrowheads = posterior growth zone; blue arrowheads = segmental stripes; brown arrowheads = parapodia; black arrowheads = ventral midline; pink arrowheads = segmental muscles; blue asterisks = expression in cells at the border between last non-amputated segment and regenerated region; brown asterisks = expression in the last non-amputated segment; pink asterisk = blood vessels; black asterisk = early expression in the ventral part of the wound epithelium, abutting the ventral nerve cord.

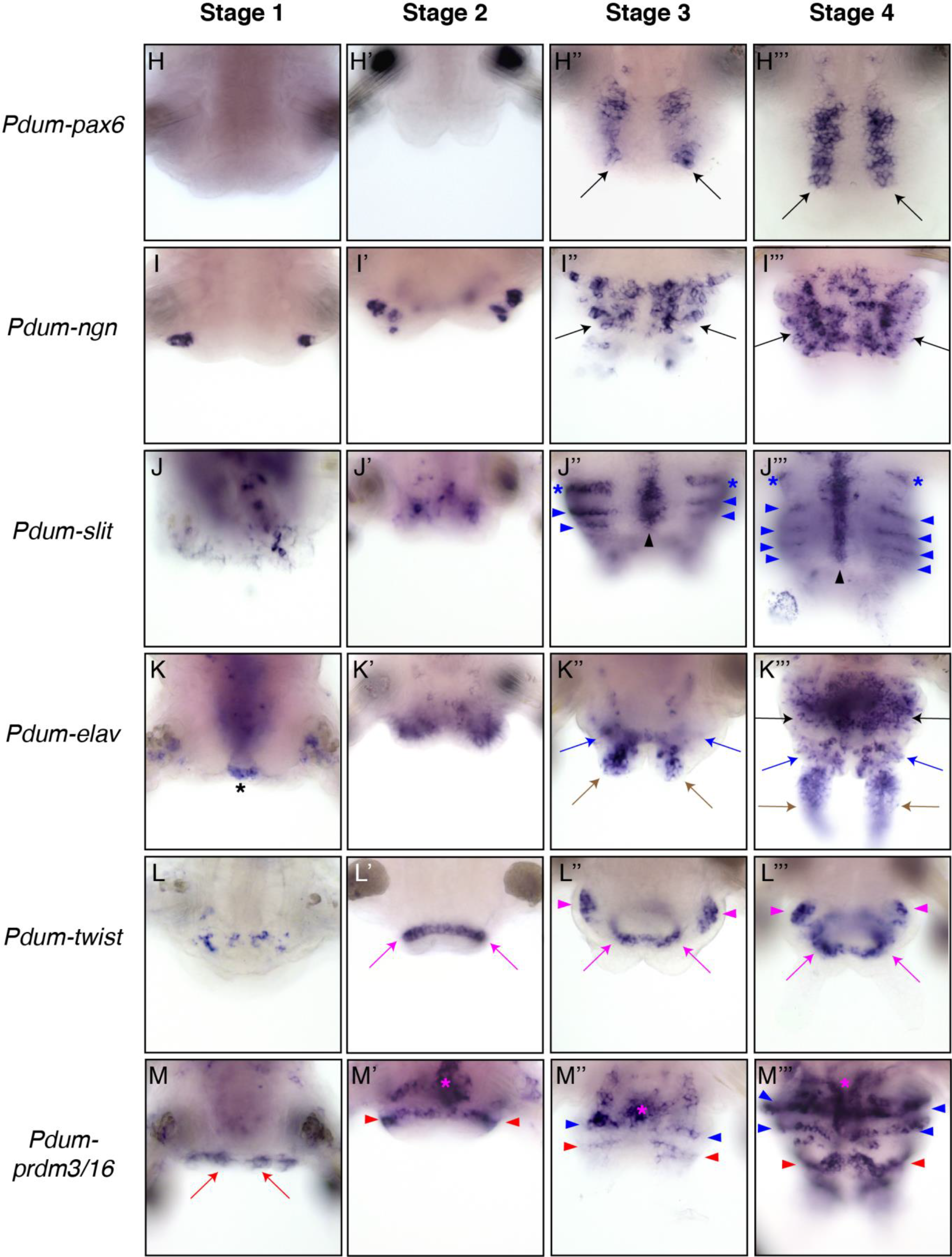

We first studied the expression during regeneration of three genes known to be expressed in the growth zone and the pygidium (*Pdum-hox3*, *Pdum-evx* and *Pdum-cdx*; Fig. 6A-C’’’; Fig. S1A-F; de Rosa et al., 2005; Gazave et al., 2013). The three genes are expressed at stage 1 in the posterior-most cells of the body, including the wound epithelium (Fig. 6A,B,C; red arrows). Their expression domains slightly extend at stage 2 (Fig. 6A’,B’,C’) and from stage 3 become very similar to those found at stage 5, with a clear expression of *Pdum-hox3* and *Pdum-evx* in a ring of cells corresponding to the growth zone and a more complex expression of *Pdum-cdx* in both the growth zone and the pygidium (Fig. 6A’’-C’’’; Fig. S2A-C; red arrowheads and blue arrows, respectively). These expression patterns are therefore suggestive of an early regeneration of the growth zone, as early as stage 3.

We next monitored segment formation and for that purpose, we studied the expression of *Pdum-engrailed* (*Pdum-en*) and *Pdum-wnt1* (Prud′homme et al., 2003), which are expressed during early or late phases of segment formation, respectively (Fig. 6D-E’’’; Fig. S1G,H). *Pdum-wnt1* is also strongly expressed in the posterior-most part of the gut and in the pygidial epidermis (Fig. 6E’-E’’’; Fig. S1H,I; green and blue arrows, respectively; Janssen et al., 2010). Both genes are expressed at stage 1 in the wound epithelium (Fig. 6D,E, red arrows). From stage 2 to stage 4, *Pdum-en* is strongly expressed in a stripe of cells located at the border between the last non-amputated segment and the regenerated region, as well as in stripes of ectodermal cells in the regenerated region (Fig. 6D’-D’’’; blue asterisks and arrowheads, respectively). From stage 2, *Pdu-wnt1* is expressed in the posteriormost part of the gut and, from stage 3, also in peripheral posterior cells, probably pygidial epidermal cells (Fig. 6E′-E’’’; green and blue arrows, respectively). In addition, at stage 4, one or two faint ectodermal stripes of *Pdum-wnt1* expression can be observed on the dorsal side of the regenerating part of the worms (Fig. S2D; blue arrowheads). At stage 5, additional stripes of *Pdum-en* and *Pdum-wnt1* expression are found (Fig. S1G,H; blue arrowheads). We conclude that segment formation starts very early during regeneration: a first segment could be specified at stage 2 and additional segments are added from stage 3 to 5, suggesting the presence of an already active growth zone from early stages.

One conspicuous feature of *P. dumerilii* segments is the presence of appendages, the parapodia. An early marker of appendage formation in *P. dumerilii* is *Pdum-dlx* which is expressed in the primordia of all body outgrowths including parapodia and cirri (Grimmel et al., 2016). At stage 1 and 2, *Pdum-dlx* is expressed in two bilateral patches of cells that correspond to the position where the anal cirri will form (Fig. 6F-F’; brown arrows) and, at later stages, in a large group of cells at the basis of the forming anal cirri (Fig. 6F’’-F’’’; brown arrows). In addition, at stage 3, we also observed two small bilateral groups of *Pdum-dlx*-expressing cells in a more anterior part of the regenerated region, which likely correspond to the anlage of the parapodia of two developing segments (Fig. 6F’’; brown arrowheads). Developing parapodia with a large number of *Pdum-dlx*-expressing cells are observed at stage 4 and 5 (Fig. 6F’’; Fig. S1J; brown arrowheads). We also analyzed the expression during regeneration of *Pdum-sfrp1/2/5* (Bastin et al., 2015), which encodes a putative secreted antagonist of Wnt signaling and which is expressed in the developing parapodia (Fig. S1K,L; brown arrowheads), as well as in ectodermal stripes (Fig. S1K,L; blue arrowheads) and in the ventral midline of the central nervous system (Fig. S1K,L; black arrowheads). At stage 1, this gene is strongly expressed in the wound epithelium (Fig. 6G, red arrows). At stage 2, there is still a weak expression in the wound epithelium, but strong expression can be observed in the gut (Fig. 6G’, green arrows; Fig. S2F), the dorsal epidermis and in the posterior part of the parapodia of the differentiated segment that abuts the amputation site (Fig. 6G’, brown asterisks; Fig. S2F). At stage 3, the two latter sites of expression are still observed while the expression in the gut fades away (Fig. 6G’’; Fig. S2F’). At stage 4 and 5, *Pdum-sfrp1/2/5* is expressed in cells of the developing parapodia (Fig. 6G’’’; brown arrowheads), epidermal cells (in particular on the dorsal side; Fig. 6G’’’; Fig. S2F’’; blue arrowheads) and ventral midline cells (Fig. 6G’’’; black arrowhead). A relatively late step of parapodia development is the formation of chaetae, the characteristic chitinous bristle of annelid appendages, whose early formation is marked by the expression of Chitin Synthase (CS)-encoding genes such as *Pdum-cs1* (Gazave et al., 2017; Zakrzewski et al., 2014). Expression of this gene is found only in the most anterior segment of the regenerated region at stage 4 and 5 (Fig. S1M; Fig. S2E, yellow arrowhead). No expression of this gene can be observed at earlier stages (not shown). Taken together, these expression data indicate that the parapodial anlagen could be specified as early as at stage 3 and that developing parapodia are present from stage 4.

We next focused on nervous system formation during regeneration. Prior work has shown that ventral nerve cord (VNC) formation during *P. dumerilii* larval and post-larval development involves the successive expressions of several categories of genes, such as *Pdum-pax6*, which has an early expression in the ventro-lateral part of the trunk ectoderm, *Pdum-neurogenin* (*Pdum-ngn*), expressed a bit later in most or all neural progenitors, *Pdum-slit*, expressed in ventral midline cells, and *Pdum-elav*, expressed in most or all differentiating neurons (Fig. 6 H-K’’’; Fig. S1N-Q; Denes et al., 2007; Simionato et al., 2008); Béhague, Kerner, Balavoine and Vervoort, unpublished observations). *Pdum-pax6* is expressed from stage 3 to 5 in two longitudinal ventro-lateral bands of cells in the anterior part of the regenerated region, which likely correspond to neurectodermal domains of the future segments (Fig. 6H-H’’’; black arrows). No expression is observed at earlier stages (Fig. 6H-H’). *Pdum-ngn* is expressed at stage 1 and 2 in a few lateral cells of unknown identity (Fig. 6I-I’). From stage 3, it starts to be expressed in many cells on the ventral side of the regenerated region, much probably neural progenitors of both the VNC and the peripheral nervous system of the developing segments, as well as in some cells of the future pygidium, in particular cells associated to the anal cirri (Fig. 6I’’-I’’’; black arrows). *Pdum-slit* is first expressed in a few scattered cells at stage 1 and 2, and from stage 3 in ventral midline cells as well as in ectodermal segmental stripes (Fig. 6J-J’’’; black and blue arrowheads, respectively). *Pdum-elav* is expressed at stage 1 in cells of the ventralmost part of the wound epithelium, in cells that are close to the amputated VNC (Fig. 6K; black asterisk). At stage 2, unlike other studied neural gene such as *Pdum-ngn* and *Pdum-pax6*, *Pdum-elav* is expressed in many cells in the ventral part of the regenerated region (Fig. 6K’). From stage 3, *Pdum-elav* expression is observed in many cells, likely the differentiating neurons of the developing segments (Fig. 6K’’’; black arrows), pygidium (Fig. 6K’’,K’’’; blue arrows) and anal cirri (Fig. 6K’’,K’’’; brown arrows). To further characterize nervous system regeneration, we performed immunolabelings with antibodies against acetylated tubulin at different stages of regeneration (Fig. 7A-A’’’’’). We observed at stage 1 and 2 the presence of bilateral nerves that extends from the VNC towards the dorsal side of the worm and that underlie the wound epithelium (Fig. 7A’,A’’; red arrows). From stage 3, a well-differentiated VNC is present in the regenerating region (white asterisk), which progressively becomes more complex and produce peripheral nerves (Fig. 7A’’’-A’’’’’; white arrowheads). To better understand when the bilateral nerves observed at stage 1 start to develop, we performed immunolabelings at different time points between amputation time and stage 1 (1dpa). We found that, while not observed at the time of amputation, these nerves are already present as early as five hours post amputation (5 hpa; Fig. S3A-A’’’; red arrows). Taken together, these data indicate that nervous system formation in the regenerated part has already started at stage 3 and follows similar steps as compared to its initial formation during development.

**Figure 7:**
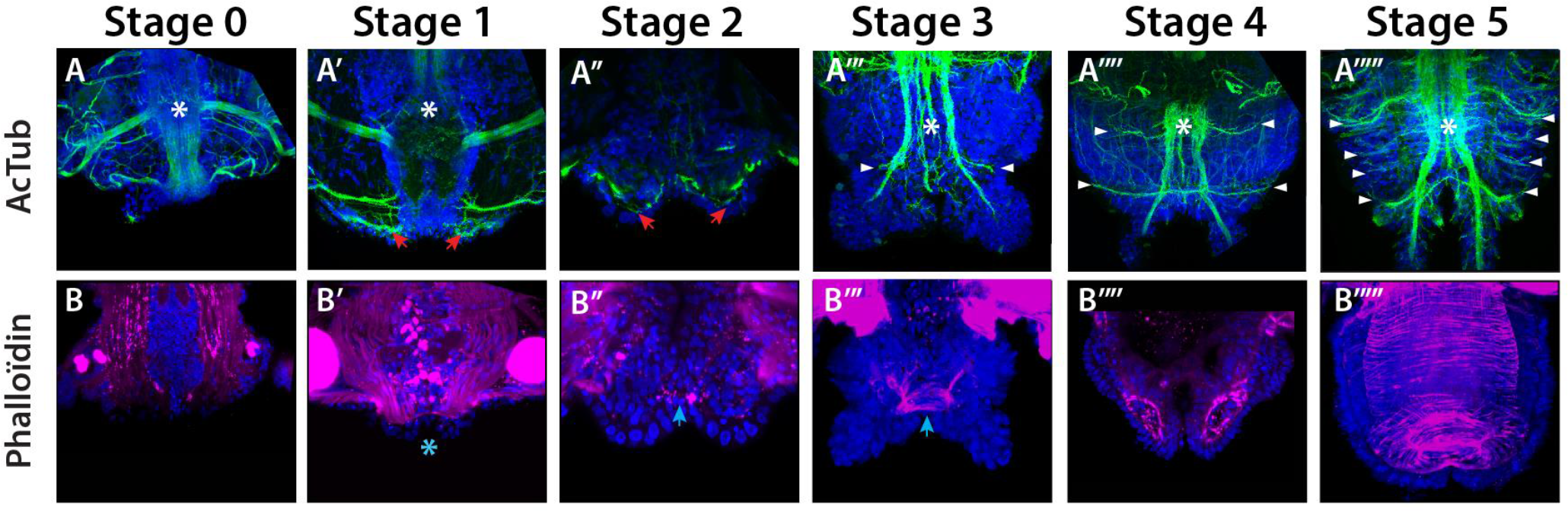
Nervous system and muscle differentiation during posterior regeneration. All pictures are ventral views (anterior is up) of posterior part of worms counterstained with Hoechst nuclear staining (in blue). (A-A’’’’’) Antibody labeling against acetylated tubulin (green) shows the axon scaffold of the ventral nerve cord (VNC, white asterisk) and peripheral nerves (white arrowheads). Stage 0 corresponds to worms that have been fixed immediately after amputation. At stages 1 and 2 bilateral nerves extend from the VNC and underlie the wound epithelium (red arrows). From stage 3 onward, peripheral nerves appear, some of which extending from the VNC in the tentacles. (B-B’’’’’) Phalloidin labeling (purple) highlights the muscle fibers. At stage 0 (B), the muscle fibers are sharp cut. One day later (stage 1, B’), the muscle fibers are contracted close to the amputation site (cyan asterisk) and at stage 2, pygidial muscles start to differentiate (B’’, cyan arrow). Later on (stage 3 to 5), segmental muscles and muscles of the gut progressively appear (B’’’-B’’’’’).

We also studied the formation during regeneration of mesodermal derivatives, namely somatic muscles and blood vessels. We used the expression of *Pdum-twist* as marker of somatic muscle development (Pfeifer et al., 2013), as this gene is expressed in the developing segmental and pygidial muscles (Fig. S1R; pink arrowheads and arrows, respectively). At stage 1, a weak expression is observed in a few scattered cells (Fig. 6L). At stage 2, *Pdum-twist* is strongly expressed in a ring of internal cells (Fig. 6L’; pink arrows) and this expression continues in the next stages, probably concerning mesodermal cells that will produce the pygidial circular muscles (Fig. 6L’’,L’’’). In addition, from stage 3 two bilateral and more anterior groups of *Pdum-twist*-expressing cells are detected (Fig. 6L’’,L’’’; Fig. S1R; pink arrowheads), which correspond to the developing somatic segmental mesoderm. To further analyze muscle formation, we performed phalloidin labelings at different stages of regeneration, which suggest that the differentiation of pygidial muscles starts at stage 2 and that of segmental muscles at stage 4 (Fig. 7B-B’’’’’). We also studied the expression of *Pdum-prdm3/16* (Vervoort et al., 2016; Kerner and Vervoort, unpublished observations) which is expressed in the developing blood vessels (Fig. S1T; pink asterisk), as well as in ectodermal segmental stripes (Fig. S1S; blue arrowheads), the growth zone (Fig. S1S,T; red arrowheads), and anal cirri (Fig. S1S; brown arrows). At stage 1, this gene is expressed in the wound epithelium (Fig. 6M, red arrows). At stage 2 and 3, *Pdum-prdm3/16* expression is observed in dorsal and ventral medial patches of cells that might correspond to precursor cells of the blood vessels (Fig. 6M’,M’’; Fig. S2G,G’; pink asterisk), as well as in the growth zone (Fig. 6M’,M’’; red arrowheads) and developing segments (Fig. 6M’,M’’; blue arrowheads). These latter expression sites are also found at later stages where a clear expression in both ventral and dorsal blood vessels is also observed (Fig. 6M’’,M’’’; Fig. S1S,T; Fig. S2G’’; pink asterisk). The specification of mesodermal structures seems therefore to start as early as at stage 2.

In conclusion, our gene expression study, summarized in figure S4, indicates that *P. dumerilii* posterior regeneration is a very fast process: at stage 3, *i.e.*, three days after amputation, a functional growth zone is present and has already produced segments in which neurogenesis and mesodermal structures formation are ongoing, and the pygidium has already undergone its differentiation. Some tissue specification, in particular mesodermal tissues, occurs even earlier, at stage 2.

### *Stem cell gene expression during* P. dumerilii *posterior regeneration*

We previously showed that homologs of several genes known to be expressed in other animals in somatic and/or germinal stem cells (hereafter named ‘stem cell genes’) were expressed in the posterior growth zone in *P. dumerilii* (Gazave et al., 2013). The posterior growth zone contains two subpopulations of putative stem cells, one superficial (probably ectodermal) and the other internal (probably mesodermal), which express different combinations of stem cell genes and are located on the ventral and dorsal side of worms, respectively (Gazave et al., 2013). Most of these genes are expressed not only in the growth zone cells, but also in a graded manner in mesodermal and/or ectodermal cells of developing segments (Gazave et al., 2013). In the present work, we studied the expression of some of these genes during regeneration. This includes a previously characterized *piwi* gene (Rebscher et al., 2007). Using available transcriptomic data (Chou et al., 2016), we identified and cloned a second *P. dumerilii piwi* homolog – the previously known gene and the newly discovered one were named *Pdum-piwiA* and *Pdum-piwiB*, respectively, based on phylogenetic analysis (Fig. S5). Expression patterns of all the selected stem cell genes at stage 5 are shown in Fig. S6. These expressions are mainly similar to those previously reported, with a few differences, in particular the fact that we did not detect *Pdum-piwiA* and *Pdum-vasa* expression in the ectodermal part of the growth zone, in contrast to what was previously observed in older worms 12 days after amputation (Gazave et al., 2013). The expression patterns of all selected stem cell genes at stages 1 to 4 of regeneration are shown in Figure 8.

At stage 1, *Pdum-piwiA* and *Pdum-piwiB* genes are expressed in cells of the differentiated segments immediately adjacent to the amputation site (Fig. 8A,B). No clear expression of *Pdum-vasa* can be detected at this stage, while *Pdum-pl10* is expressed in many cells of the posterior part of the differentiated segment abutting the amputation plane (Fig. 8C,D). At stage 2, *Pdum-piwiA* and *Pdum-vasa* are expressed in bilateral groups of internal cells in the regenerated region (Fig. 8A’,C’). *Pdum-piwiB* and *Pdum-pl10* are expressed in most cells of this region (Fig. 8B’, D’), the latter being also expressed in superficial cells on the ventral side (not shown). From stage 3, the four genes are expressed in the mesodermal growth zone and mesoderm of the developing segments (Fig. 8A’’-D’’’; red arrowheads). *Pdum-pl10* is in addition strongly expressed in cells at the basis of the anal cirri (Fig. 8D’’-D’’’; brown arrows) and in the ectodermal part of the growth zone (Fig. S2H; red arrowheads). At stage 5, *Pdum-piwiA*, *Pdum-piwiB*, *Pdum-vasa*, and *Pdum-pl10* are expressed in the mesodermal part of the growth zone (Fig. S6A-D; red arrowheads), mesodermal cells of the developing segments, and more or less strongly in cells at the basis of the anal cirri (Fig. S6A-D; brown arrows).

**Figure 8:**
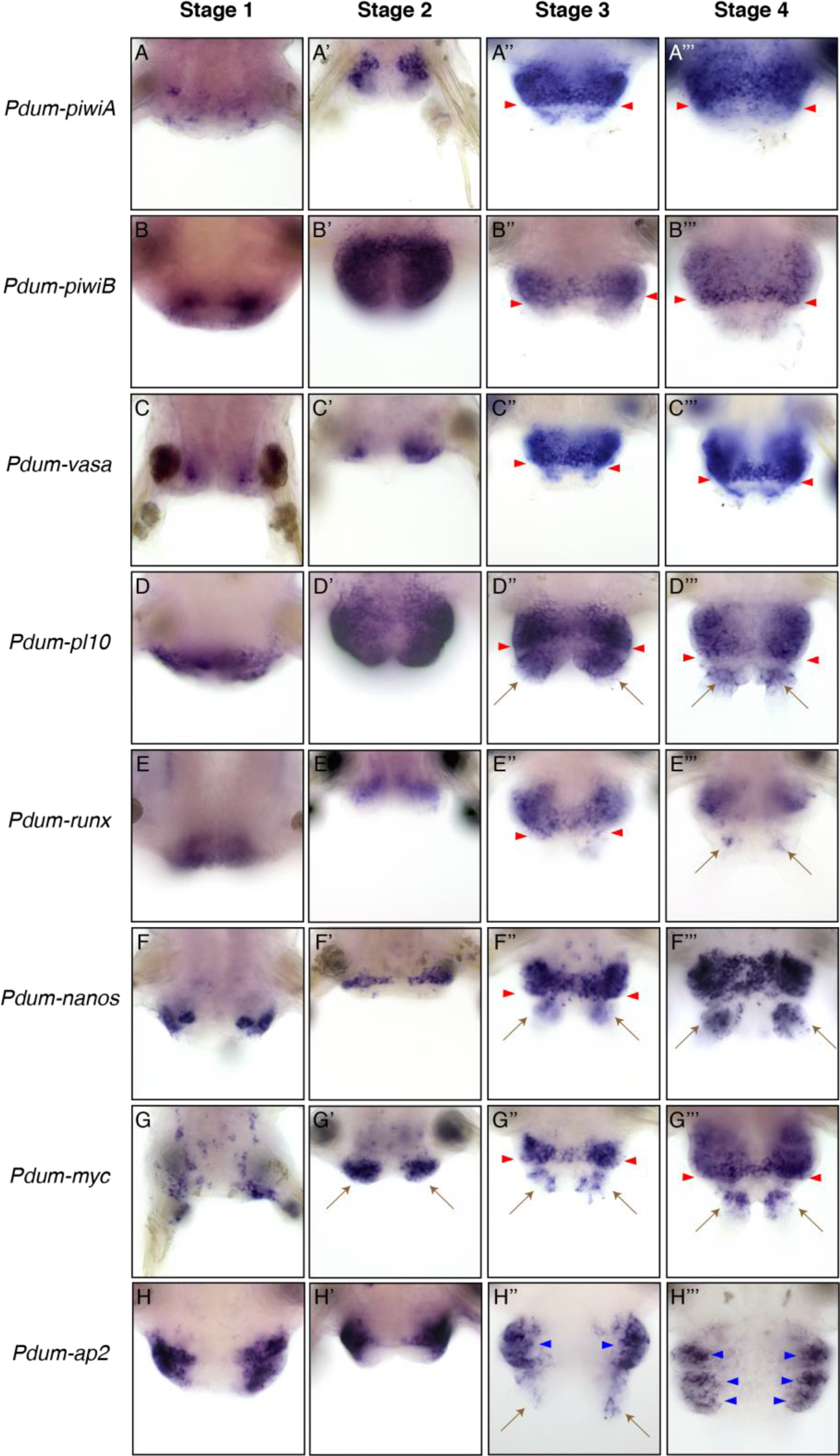
Expression of ‘stem cells genes’ during posterior regeneration. Whole-mount *in situ* hybridizations (WMISH) for the genes whose name is indicated are shown for four posterior regeneration stages (stage 1 to stage 4). Stage 5 WMISH images are shown in Fig. S5. All panels are ventral views (anterior is up). brown arrows = anal cirri; red arrowheads = posterior growth zone; blue arrowheads = segmental stripes.

A stage 1, *Pdum-runx* is weakly expressed in the ventral posteriormost part of the last differentiated segment (Fig. 8E). Weak expression in internal cells in the regenerated region is observed at stage 2 (Fig. 8E’). At stage 3, it is expressed in the mesodermal growth zone and the developing mesoderm (Fig. 8E’’; red arrowheads). The expression in the growth zone fades away at stage 4 and a weak expression in the anal cirri can be observed (Fig. 8E’’’; brown arrows). At stage 5, *Pdum-runx* is weakly expressed in the developing mesoderm (Fig. S6E). We did not detect clear expression in the mesodermal growth zone in contrast to what has been reported in older worms (Gazave et al., 2013). *Pdum-nanos* is expressed in two lateral patches of superficial cells in the differentiated segment close to the amputation at stage 1 (Fig. 8F). Its expression extends ventrally at the border between the differentiated segment and the regenerated region (Fig. 8F’). At stage 3 and 4, *Pdum-nanos* expression is similar to that seen at stage 5 (Fig. S6F), with strong expression in the ectodermal growth zone (Fig. 8F’’-F’’’; red arrowheads), developing nervous system and anal cirri (Fig. 8F’’-F’’’; brown arrows). At stage 1, *Pdum-myc* is expressed in many scattered cells in the last differentiated segment (Fig. 8G). Strong expression of the gene is observed at stage 2 in two posterior patches of cells located where the anal cirri will form (Fig. 8G’; brown arrows). At stage 3 and 4, *Pdum-myc* is expressed in both ectodermal and mesodermal parts of the growth zone (Fig. 8’’,G’’’G; Fig. S2I; red arrowheads), developing segments and anal cirri (Fig. 8G’’,G’’’; Fig. S2I; brown arrows). Expression is detected at stage 5 in the mesodermal and ectodermal parts of the growth zone, in the developing segmental mesoderm and ectoderm, and in anal cirri (Fig. S6G; red arrowheads and brown arrows, respectively). At stage 1, *Pdum-ap2* is strongly expressed in two dorso-lateral patches of cells on the ventral side of the last differentiated segment (Fig. 8H) and in the dorsal part of the wound epithelium (Fig. S2J). Strong dorso-lateral and dorsal expression can also be observed at later stages and additional expression in the developing anal cirri is also observed (Fig. 8H’-H’’’; Fig. S2J’-J’’’; blue arrowheads, red arrowheads and brown arrows, respectively). At stage 5, *Pdum-ap2* is expressed in dorso-lateral stripes of epidermal cells (Fig. S6H, blue asterisks and arrowheads) and in the ectodermal part of the growth zone (Fig. S6I, red arrowheads).

All the studied stem cell genes are therefore expressed during *P. dumerilii* posterior regeneration and several of them are broadly expressed during early stages of the process (stage 2 and even stage 1 for some genes). As these genes are usually expressed in proliferating cells in other models and during *P. dumerilii* growth (Gazave et al., 2013), we next studied cell proliferation at different stages of regeneration.

### *Cell proliferation during* P. dumerilii *posterior regeneration*

As a first approach to identify proliferating cells, we studied the expression of three previously characterized cell cycle genes, *Pdum-cycB1*, *Pdum-cycB3* and *Pdum-pcna* (Demilly et al., 2013; Gazave et al., 2013). *Pdum-cycB1* and *Pdum-cycB3* have similar expression patterns throughout the regeneration process (Fig. 9A-B’’’; Fig. S6J,K). At stage 1, the two genes are expressed in few cells located close to the wounded epidermis (Fig. 9A,B). At stage 2, strong expressions are detected in many cells of the regenerated region (Fig. 9A’,B’). Strong expressions are also observed at stage 3, 4 and 5 in the mesodermal part of the growth zone, the future mesoderm of the developing segments and at the basis of the anal cirri (Fig. 9A’’, A’’’, B’’, B’’’; Fig. S6J,K; red arrowheads and brown arrows). At stage 1, *Pdum-pcna* is strongly expressed in two large lateral patches of cells at the amputation site, that include cells of the wound epithelium (Fig. 9C). At later stages, *Pdum-pcna* expression is similar to that of the two *cyclin* genes, while being stronger and probably concerning more cells than the other cell cycle genes (Fig. 9C’-C’’’; Fig. S6L). In addition, strong expression of *Pdum-pcna* is observed in the ectoderm, including the ectodermal part of the growth zone (Fig. S2K-K’’; Fig. S6M), in contrast with *Pdum-cycB1* and *Pdum-cycB3*. We also performed immunolabelings with a cross-species reactive anti-PCNA antibody, previously used in another annelid (Niwa et al., 2013). PCNA immunoreactivity is found in cells at all stages of the regeneration process (Fig. S7A-E’), including cells of the intestine (Fig. S7C, D) and the growth zone at late stages (Fig. S7D’-E’, red arrowheads), albeit in much lower numbers of cells as compared to WMISH of *Pdum-pcna*. We do not know whether this difference reflects real differences between RNA and protein distributions or might have some other (technical for example) reasons.

**Figure 9:**
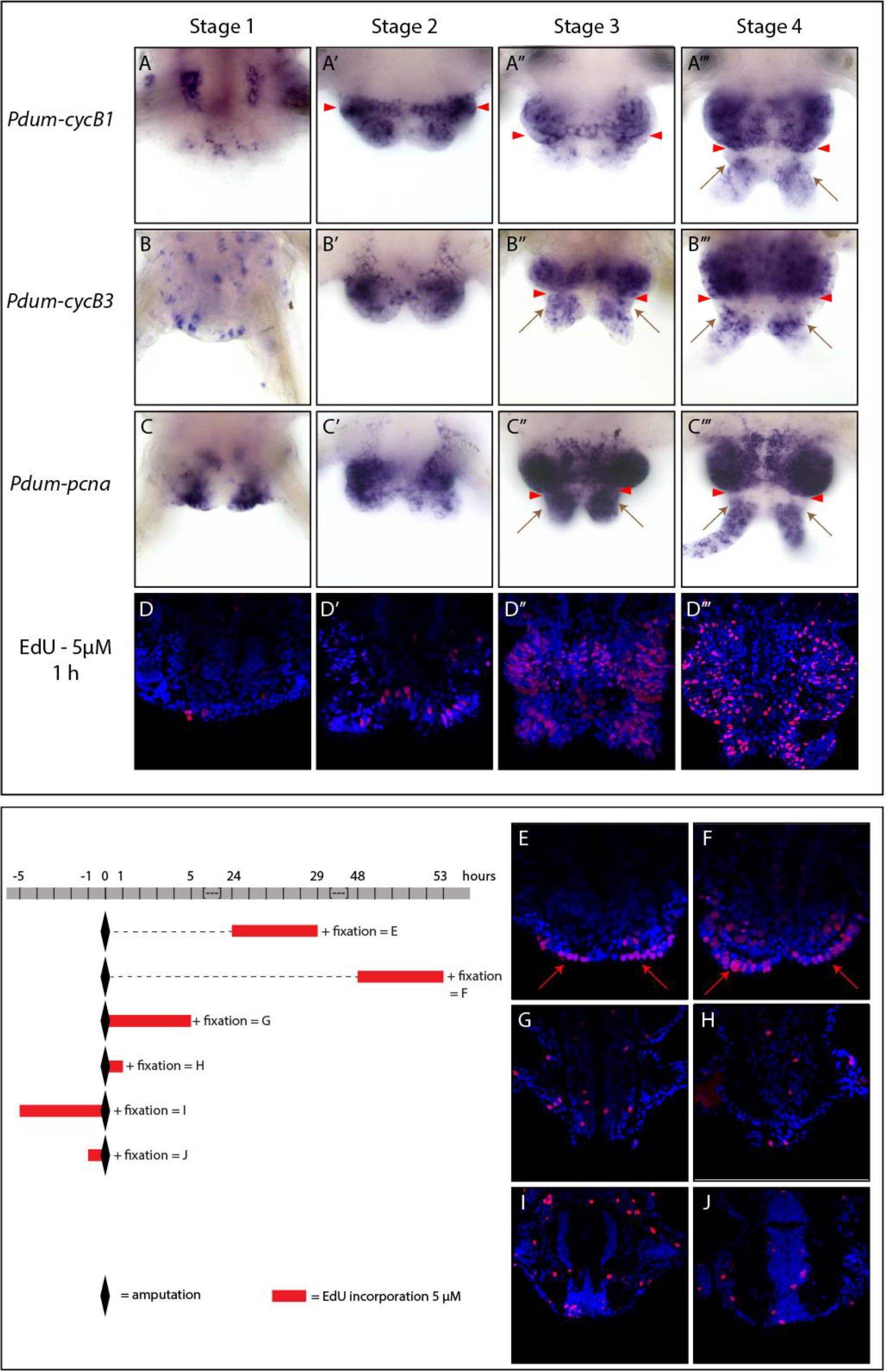
Cell proliferation during posterior regeneration. (A-C’’’): Whole-mount *in situ* hybridizations (WMISH) for three cell cycle genes (*Pdum-cycB1*, *Pdum-cycB3* and *Pdum-pcna*) are shown for four posterior regeneration stages (stage 1 to stage 4). All images are ventral views, anterior is up. (D-J): Posterior parts of worms (anterior is up) showing EdU labeling (red) and Hoechst nuclear staining (blue). (D-D’’’): 1hour of EdU incorporation made at different stages of posterior regeneration. (E-J): 1-and 5-hour EdU incorporations made at different time points as described in the scheme in the left part of the lower panel. The gray bar represents a time scale, 0 being the time point when amputations were made (indicated by black diamonds). Red bars indicate when and for how long EdU incorporations were performed. Red arrowheads: posterior growth zone; brown arrows: anal cirri; red arrows: wound epithelium.

To further evaluate cell proliferation during posterior regeneration, we performed EdU (5-ethynyl-2’-deoxyuridine) incorporations (to label cells that are in S phase) and anti-phospho-Histone H3 (anti-PH3) immunolabelings (to label cells in M phase). In a first set of experiments, we incorporated EdU for one hour at the different stages of regeneration, followed by immediate fixation (Fig. 9D-D’’’; stage 5 not shown as EdU+ positive cell distribution is identical to that previously described during posterior elongation; Gazave et al., 2013). At stage 1 and 2, very few cells are labeled near the amputation site (stage 1; Fig. 9D) and in the regenerated region (stage 2; Fig. 9D’). At later stages, much more labeled cells are detected in the regenerated part, mainly in its anterior part that will give rise to segments and in the anal cirri (Fig. 9D’’-D’’’; not shown for stage 5). Similarly, while only very few cells are labeled with anti-PH3 at stage 1 and 2, more labeled cells are found at the later stages, much less numerous than the EdU+ cells, but with a comparable distribution (Fig. S7F-J). The expression of cell cycle genes and the distribution of PH3+ or EdU+ cells point out strong proliferation in the regenerated region from stage 3 onward. There are however some discrepancies at the earlier stages, in particular stage 2, as at this stage wide expressions of *Pdum-cycB1*, *Pdum-cycB3* and *Pdum-pcna* and only few EdU+ or PH3+ cells are observed. In order to evaluate more precisely cell proliferation during early steps of regeneration, we performed longer EdU incorporations (5 hours) and found much more labeled cells than with one-hour incorporations, located mainly in the wound epithelium at stage 1 and, at stage 2, in both ectodermal and mesodermal cells of the regenerated region and in the differentiated segment adjacent to the amputation site (Fig. 9E-F, respectively). One and five hours EdU incorporations performed immediately before and after amputation reveal a very low level of basal cell proliferation, as very few EdU+ positive cells are observed (Fig. 9G-J). Consequently, the pattern of proliferation observed from stage 1 onward is likely a response to amputation.

Taken together, our data indicate that posterior regeneration in *P. dumerilii* involves high levels of cell proliferation, raising the possibility that cell proliferation may be required for regeneration to proceed in a normal way. To test this hypothesis, we treated worms with widely-used anti-proliferative agents, hydroxyurea (HU) and nocodazole, and analyzed their effects on regeneration. While completely blocking regeneration, nocodazole was very toxic, even at low concentration (0.01μM), and caused the death of most worms during treatment (not shown), precluding us from drawing from these experiments firm conclusions about the role of cell proliferation during regeneration. HU in contrast appeared to be much more harmless to worms: in a first concentration range test experiment, we put worms immediately after amputation in three different HU concentrations (10mM, 20mM and 50mM) for five days and scored the worms every day for the stage that has been reached (Fig. 10A). No worm death was observed and with the three used concentrations regeneration was much delayed as compared to controls: at 5dpa, while control worms were at stage 5, worms treated with HU were mostly at stage 2 and no stage 4 or 5 was observed. While HU-treated worms reached stage 1 at 1dpa like controls, significant regeneration delays started to be detected at 2dpa for the 50mM HU condition. At 5dpa, worms treated with 50mM HU tended to regress to previous stages, which could reflect toxic effects of HU at this concentration. We therefore chose 20mM concentration for further experiments as in this condition these putative toxic effects were not observed. In addition, worms treated with this concentration present less variability in the stage that has been reached than the worms treated with the lowest concentration (Fig. 10A). To confirm that HU did indeed block cell proliferation in our experiments, we used an EdU pulse and chase experimental paradigm: one-hour EdU incorporation was performed at 3dpa to label cells in S phase and followed by a two-day chase either in sea water (controls) or sea water with 20mM HU (Fig. 10B). As expected, in control worms, many EdU+ positive nuclei were observed and many of them showed stippled EdU labelings that are typical of cells produced by cell divisions having occurred since incorporation (Fig. 10B1; *e.g.*, Demilly et al., 2013). In contrast, in HU treated worms, a reduced number of labeled nuclei were observed and all nuclei showed a strong and homogenous labeling, indicating that cells did not divide in the presence of HU (Fig. 10B2). To better understand how regeneration proceeds in presence of HU, we did HU treatment from 0dpa to 5dpa, fixed the treated worms at 5dpa and performed WMISH for some of the genes whose expression was studied during normal regeneration (Fig. S8). All the analyzed genes showed expressions at 5dpa in HU treated worms similar to those of stage 2 in non-treated worms, further indicating that regeneration is blocked in presence of HU.

**Figure 10:**
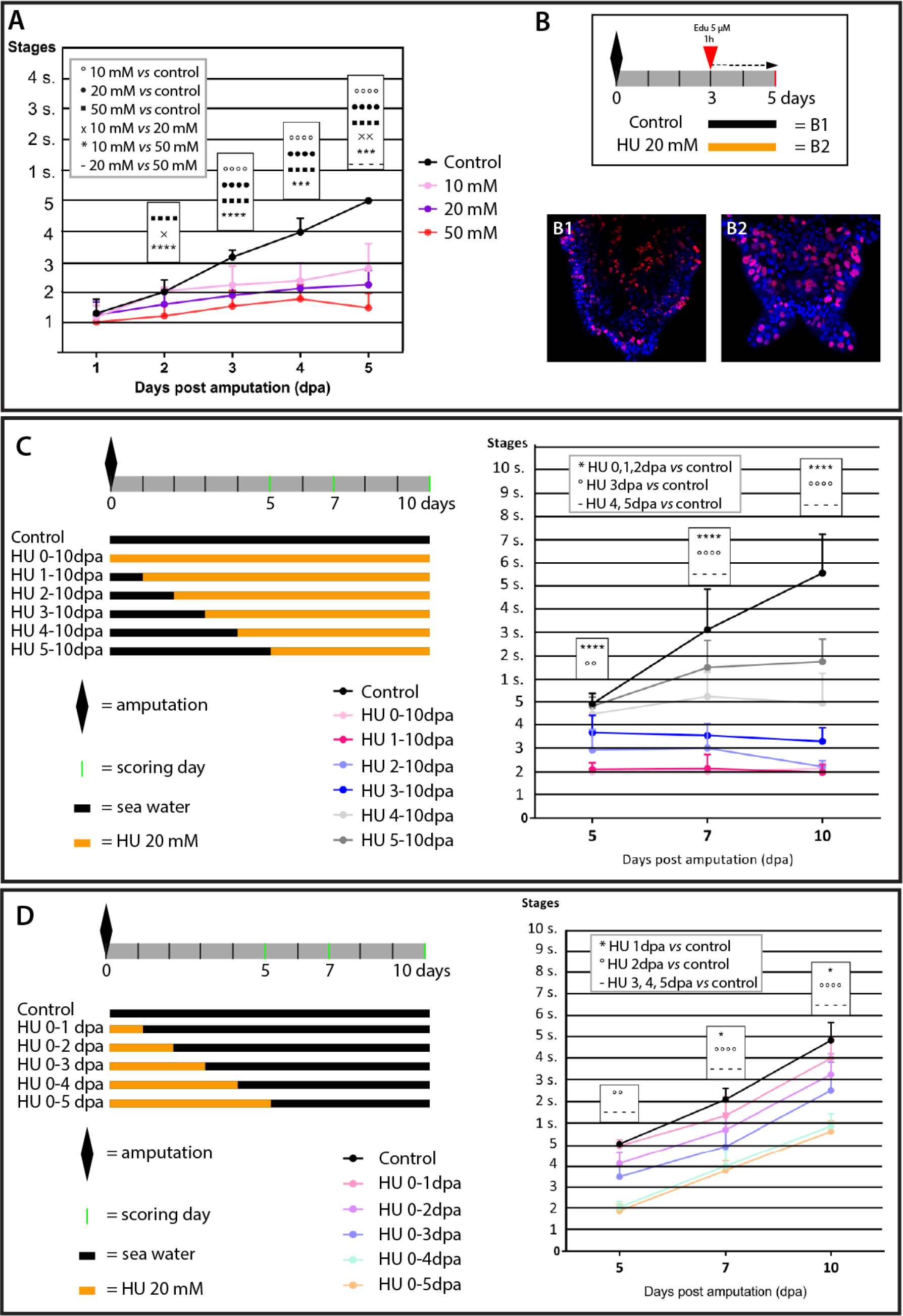
The anti-proliferative agent hydroxyurea (HU) impairs posterior regeneration in a reversible manner. (A) Graphic representation of the stages reached by control worms and worms treated with three different concentrations of HU (n=18 per HU conditions, n=14 for control condition) every day during five days. HU-treated worms showed a significantly delayed regeneration compared to that of controls. (B) Schematic representation of the experimental design used to confirm HU anti-proliferative effect: 3dpa worms were incubated 1hour with 5μM EdU and chased in normal sea water (controls; B1) or sea water with 20mM HU (B2) for 2 days (until 5dpa) before fixation. (B1) and (B2): EdU labelings of control worms (B1) and HU-treated worms (B2) showing that HU stops cell divisions. In (C) and (D), grey bar represents the time line of the experiments and green bars the scoring days. Orange bars represent time periods during which worms were incubated in HU 20mM in sea water, black bars time period of incubation in normal sea water (without HU). (C) On the left, schematic representation of HU treatments. On the right, graphic representation of the stages reached by the worms in the different conditions (n=12 per HU condition, n=16 for control condition). Significantly delayed or blocked regeneration was observed. In all conditions, worms were nevertheless able to reach stage 2. (D) On the left, schematic representation of HU treatments. Stages reached by the worms in the control and five experimental conditions (n=12 per condition). On the right, graphic representation of the stages reached by the worms in the different conditions. Statistics for (A), (C) and (D): 2-way ANOVA on repeated measures using Dunnett correction were done. * p<0.05; ** p<0.01; *** p<0.001; **** p<0.0001. Error bar: SD

In order to identify at which stages of regeneration cell proliferation is required, we next performed HU treatment starting at different time points after amputation (from 0dpa to 5dpa) and pursued treatment until 10dpa, scoring the stages reached by worms at 5dpa, 7dpa and 10dpa (Fig. 10C). Worms treated from 0 and 1dpa reached stage 2 but did not progress beyond this stage. Worms treated from 2dpa were between stage 3 and 4 at 5dpa and 7dpa, but their regenerated region subsequently tended to regress, so at 10dpa most of these worms present a stage 2 morphology. Treatment from 3dpa did not allow the worms to go beyond stage 4 and some morphological regressions were also observed in these worms after day seven. Worms treated from 4 and 5dpa were able to reach stage 5 at 5dpa, as the control group, but subsequent addition of segments was severely impaired (Fig. 10C). Cell proliferation is therefore required to reach stage 3 and for further progression of regeneration and segment addition. We also tested if HU treatment was reversible by starting HU treatment immediately after amputation (0dpa) and stopping it at different time points (1 to 5dpa). As in the previous experiment, worms were scored at 5dpa, 7dpa and 10dpa (Fig. 10D). In all conditions, HU treatment was reversible: after removal of HU, regeneration started again and followed a normal path in terms of stages and timing of these stages. Interestingly, worms treated from 0 to 1dpa showed delayed segment addition and those treated from 0 to 2dpa delayed regeneration. It suggests that cell proliferation, while not required to reach stage 2, is nevertheless important during the two first days after amputation for further progression of the process.

Taken together those experiments show that intense cell proliferation occurs during *P. dumerilii* regeneration and is mandatory for regeneration to proceed beyond stage 2 of the process.

#### *Insight into the cellular origin of the* P. dumerilii *regenerated region*

Identifying the origin of the cells of the regenerated region is an important aspect of the understanding of a regeneration process. Pulse and chase experiments have been widely used to indirectly tackle this question (*e.g.*, de Jong and Seaver, 2017) and we designed two such experiments to study *P. dumerilii* posterior regeneration. In a first set of experiment, we incorporated EdU during five hours before amputation and put the worms after amputation in normal sea water until 3dpa or 5dpa (Fig. S9). At 3dpa, a few labeled cells were found in the regenerated region, but only in internal tissues (Fig. S9A-A’). At 5dpa, many labeled cells are found in the lining of the regenerating gut and a few ones (very weakly labeled) in other internal tissues, likely mesodermal derivatives such as muscles (Fig. S9B-B’). No EdU+ cells were found in the epidermis neither at 3dpa nor at 5dpa. We next performed a second slightly more complicated pulse and chase experiment (Fig. 11A). A five-hour EdU pulse was performed one day after amputation, which labeled a significant number of cells both at the amputation site and in many differentiated segments (Fig. 11B-B’). Worms were then allowed to regenerate during two days in sea water (48h chase; 3dpa stage). As expected, the regenerated region contained many EdU labeled cells (Fig. 11C), but we cannot ascertain whether these cells derive from cells at the amputation site, or from differentiated segments located further away, or both. We therefore performed a second amputation (Fig. 11A) using three different amputation planes, the same plane than the first amputation (therefore we only removed the regenerated region; referred to as condition 1 below), one segment more anterior (removal of the regenerated region and the segment abutting the first amputation plane; condition 2), or eight segments more anterior (removal of the regenerated region and the eight most posterior segments; condition 3). After this second amputation, worms were allowed to regenerate three days in sea water (72h chase; 3dpa stage) before EdU labeling analysis. In condition 1, a large number of EdU+ cells were observed and both internal and superficial (prospective epidermis) labeled cells were found (Fig. 11D-D’). Much fewer cells were EdU+ in conditions 2 and 3 and no labeled cells were found in the epidermal cell layer of the regenerated region (Fig. 11E-E’ and F-F’, respectively). It therefore suggests that the exclusive source of ectodermal cells of the regenerated region is the immediately abutting segment.

**Figure 11:**
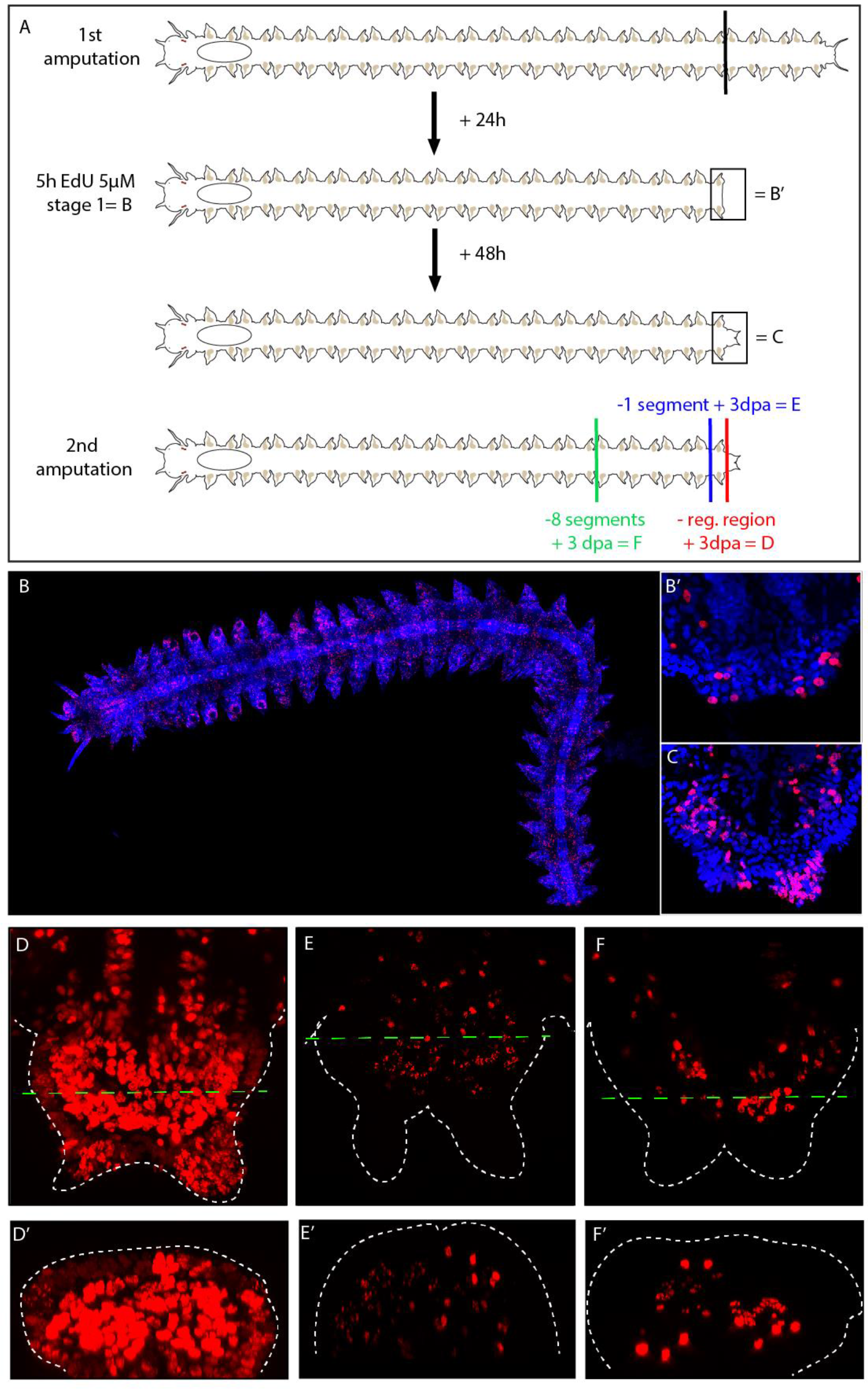
EdU pulse and chase experiments point to mostly local source of cells of the regenerated region. (A) Schematic representation of the experimental design. Corresponding pictures in the lower panel are indicated. In brief, a first amputation was performed (amputation position shown by a black bar) and 24h after this amputation EdU was incorporated for 5 hours (in sea water with 5μM EdU). Some worms were imaged (shown in B-B’). For the other worms, this incorporation was followed by a 48h chase in normal sea water. Some worms were fixed and imaged (shown in C) and the others were subsequently amputated a second time. Three different amputation planes were used (green, blue and red bars). The worms were allowed to regenerate for three days (3dpa) in normal sea water, fixed and imaged (shown in D-F’). (B-C) EdU labeling (red) and Hoechst nuclear staining (blue). (B) Whole worm EdU labeling with 1h EdU labeling 24h after amputation (anterior is on the left and posterior is down right). Many EdU+ cells are found throughout the body. (B’) Higher-magnification view of the posterior part of the worm shown in (B), showing the presence of EdU+ cells in the regenerating region. (C) Posterior part of a worm after a 48h chase in normal sea water. Many EdU+ cells are found in the regenerating region. In (B’) and (C), anterior is up. (D-F’) Posterior part of worms 3 days after the second amputation made at three different positions as schematized in (A). Anterior is up. Green dotted lines in (D), (E) and (F) indicated the position where the virtual cross-sections shown in (D’), (E’) and (F’) were done. Dorsal is up. Dotted white lines represent the shape of the worms.

A popular hypothesis is that regeneration in annelids may rely on the presence in all or most segments of neoblast-like cells, *i.e.*, stem cells expressing genes of the GMP signature (stem cell genes such as *piwi*, *myc* and *vasa*). These cells would be activated by the amputation, migrate to the amputation site and produce a part or the totality of the regenerated region (*e.g.*, Bely, 2014; de Jong and Seaver, 2017; Özpolat and Bely, 2016). There is no experimental evidence for the presence of such cells in the segments of *P. dumerilii*. However, a population of cells expressing the aforementioned stem cell genes does indeed exist at least during early post-larval stages of *P. dumerilii*: the primordial germ cells (PGCs; Rebscher et al., 2007). Four PGCs are indeed produced during embryonic development by the blastomeres that also generate the larval mesodermal progenitors and the mesodermal part of the growth zone (Özpolat et al., 2017). While located in the posterior part of the body adjacent to the growth zone in late larval stages (until four days after fertilization, dpf), PGCs then migrate anteriorly to join a region immediately posterior to the pharynx, where these cells will start to proliferate to produce germline cells (Rebscher et al., 2007). At least during their migration, these cells express most of the stem cell genes also expressed by the posterior growth zone cells (Gazave et al., 2013). An appealing hypothesis would therefore be that PGCs could in fact have both germinal and somatic potentialities, producing germ cells during normal growth and sexual maturation, but being also able to contribute to posterior regeneration if amputations or injuries occur. We tested this hypothesis, by taking benefit of a previously established procedure to label these cells based on the fact that they are quiescent during embryonic and larval development (Özpolat et al., 2017; Rebscher et al., 2012). Short EdU incorporations between 5 to 7 hours post fertilization (hpf) allow to label PGCs and many other cells. However, as most of these other cells subsequently divide a lot, after a chase until 48hpf or 72hpf, in the trunk of the larva only the four PGCs show a strong and homogenous EdU labeling (Rebscher et al., 2012). We did the same incorporation, but performed a much longer chase until worms were one-month old (Fig. 12A). We found many EdU+ cells in these worms and in particular a large number of cells in one segment posterior to the pharynx (dotted line square in Fig. 12A), much likely the progeny of the PGCs. In addition, many labeled cells were also found in the head, in agreement with previously published data showing the presence of EdU+ cells in the future head region at 72hpf (Rebscher et al., 2012), and unexpectedly also in the posterior growth zone and pygidium (Fig. 12A). To better understand this result and confirm that PGCs were labeled in this experiment, we did again EdU pulse at 5-7hpf and made chase until 3dpf, 6dpf, 10dpf and 14dpf. At 3dpf and 6dpf (Fig. 12C-D’), two groups of EdU+ cells were observed, most likely corresponding to PGCs and posterior growth zone cells (Fig. 12C’-D’; arrowheads and arrows, respectively). At 10dpf and 14dpf (Fig. 12E-F’), labeled cells at the level of the posterior growth zone and pygidium were still observed (arrows in Fig. 12E′,F’). Strongly labeled cells were also observed in more anterior position, which correspond to the migrating PGCs (Fig. 12E,F; arrowheads). We next took the 1-month worms in which PGC progeny is EdU+, amputated these worms posterior to the segment containing these cells and let the worms regenerate for three days (Fig. 12B). Strikingly, the regenerated region was almost devoid of EdU+ cells, indicating that PGC progeny does not significantly contribute to regeneration, at least in this experimental design.

**Figure 12:**
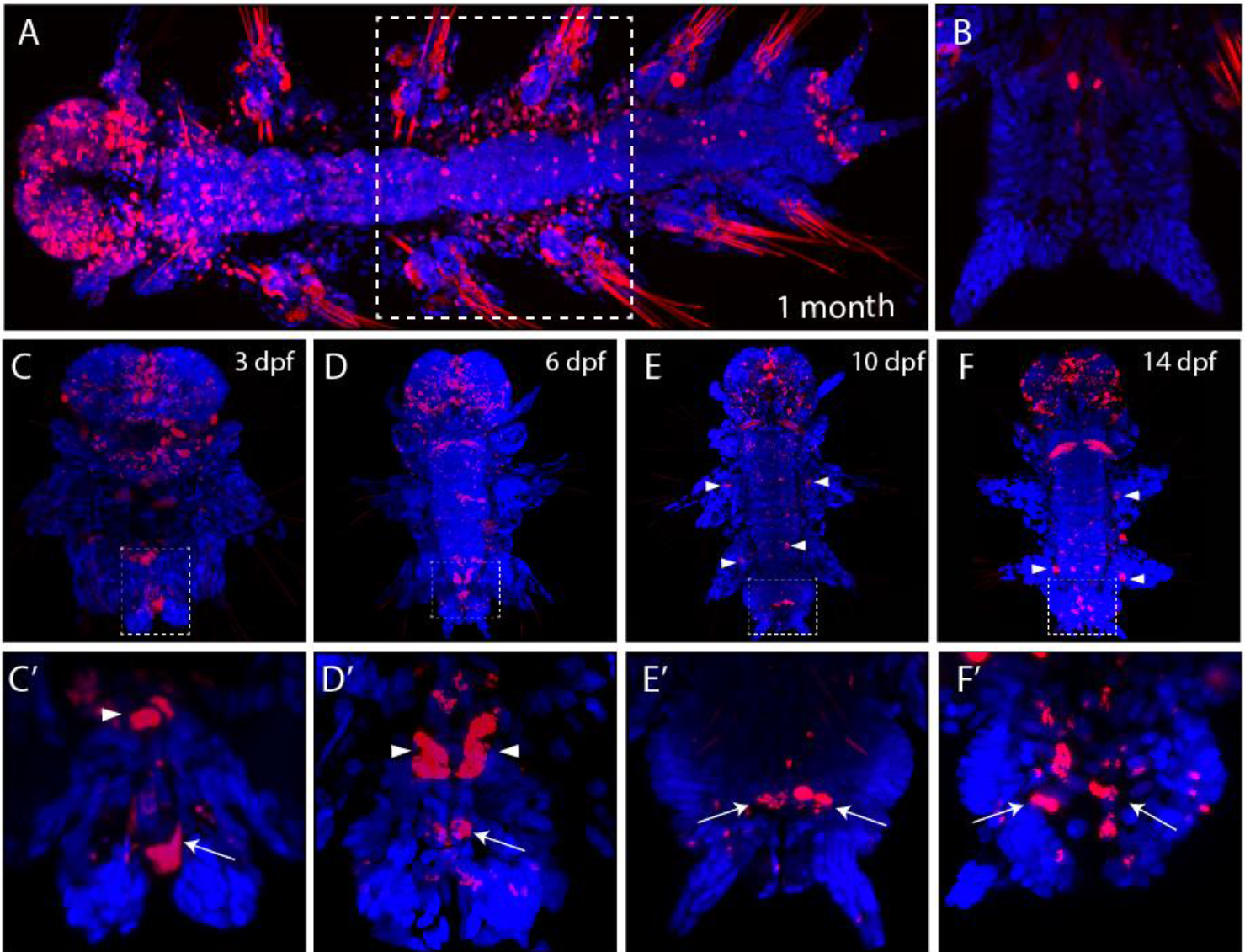
Primordial germ cells (PGCs) do not contribute to posterior regeneration. EdU incorporation was performed at 5 to 7 hours post fertilization and was followed by chase in normal sea water until 1 month (A), 3 days post fertilization (3dpf; C,C’), 6dpf (D,D’), 10dpf (E,E’), or 14dpf (F,F’). EdU labeling (in red) and Hoechst nuclear staining (in blue). In (A) anterior is on the left. In all other images, anterior is up. (C’), (D’), (E’) and (F’) are higher magnification of the posterior part of the worms shown in (C), (D), (E) and (F), respectively. White arrowheads point to PGCs and white arrows to cells of the growth zone. The dotted line square in (A) delineate the segment that contains the PGC progeny. (B) Posterior part of a 1-month old worm (in which PGC daughter cells are EdU+) three days after amputation. Almost no EdU+ cells can be found, indicating that PGC progeny does not significantly contribute to posterior regeneration.

Taken together, our data strongly suggest that cells of the regenerated region have two sources. Most cells, and notably all epidermal cells, originate from the segment immediately abutting the amputation plane, therefore pointing out a mostly local origin of the cells of the regenerated region. Some cells of the regenerated region, mostly cells of the gut (but no epidermal cells) derive from cells that were proliferative before amputation. Finally, our results do not support the hypothesis that PGCs may be a source for cells of the regenerated region.

## DISCUSSION

Many animals can regenerate complex organs or body parts after an injury or an amputation. This injury-induced regeneration ability, which extends far beyond the homeostatic repair of tissues such as epidermis and intestine epithelia found in most species, is nevertheless very widespread in metazoans (Bely and Nyberg, 2009; Grillo et al., 2016). Highly regenerative species are indeed found in most phyla, both non-bilaterians (such as cnidarians, ctenophores and sponges) and bilaterians including for example flatworms, annelids, arthropods, and chordates (Bely and Nyberg, 2009; Grillo et al., 2016). Regeneration has been the focus of intense research since the pioneering work of Abraham Trembley, Simon Pallas and Lazzaro Spallanzani in the 18^th^ century (reviewed in Sánchez Alvarado, 2000). Despite this long-standing interest and the extensive study of regeneration in species such as salamanders and planarians, fundamental questions about regeneration, in particular concerning its evolution in animals, remain still unanswered. It has been advocated that solving these questions requires to study new models (or « research organisms » using the term coined by A. Sanchez Alvarado) amenable to molecular, cellular and functional analyses (Grillo et al., 2016; Sánchez Alvarado, 2018). The Errantia annelid *P. dumerilii*, which possesses extensive regenerative abilities, appears to be a particularly appropriate model to study regeneration and its evolution, thanks to its belonging to a slow-evolving lineage and the recent establishment of sophisticated molecular and genetic tools to study its development (Raible and Tessmar-Raible, 2014; Williams and Jékely, 2016; Zantke et al., 2014). In this article, we present a detailed characterization of posterior body regeneration in *P. dumerilii*, which provides the foundation for future mechanistic and comparative studies of regeneration in this species.

### *Stages and timeline of* P. dumerilii *posterior regeneration*

Amputation of the posterior part of the body leads to the removal of the posteriormost part of the body (the pygidium), the growth zone (responsible for the addition of new segments during the posterior growth of the animal) and several segments. *P. dumerilii* worms are able to regenerate both the differentiated structures of the pygidium and the stem cells of the growth zone which in turn allows the formation of new segments that replace the amputated ones. The whole process can therefore be subdivided into two different conceptual phases, regeneration *per se*, *i.e.*, the restoration of the pygidium and growth zone, and post-regenerative posterior growth, *i.e.*, the formation of segments by the regenerated growth zone (Gazave et al., 2013). Based on morphological observations, we defined five regeneration stages that precede the appearance of well visible segments produced through post-regenerative posterior growth (Figs. 1-3). As far as homogenous worms in size and age are considered, the timeline of these stages appears highly reproducible, while the subsequent progressive addition of morphologically distinguishable segments is more variable. Our in depth characterization of the different regeneration stages, using the expression of many genes involved in cell and tissue patterning and differentiation, indicates that regeneration *per se* and post-regenerative posterior growth are largely overlapping (Fig. 6). Indeed, first signs of pygidium and growth zone restoration appear at stage 2 (*i.e.*, two days post amputation; 2dpa) and pygidium differentiation continues during the three next stages as judged by the progressive growth of the anal cirri and the progressive differentiation of the pygidial nervous system and muscles. The growth zone is regenerated between stage 2 and 3 and is undoubtedly functional as early as stage 3 (3dpa), as shown by the addition of segment primordia at this stage (Fig. 6). In stage 4 and 5, while still not visible at the morphological level, there are clearly developing segments with differentiating mesodermal and ectodermal derivatives. *P. dumerilii* posterior regeneration is therefore a very fast process and post-regenerative posterior growth in fact starts only three days after amputation. The timeline of the process roughly corresponds to that found in other annelids, for example the other Errantia-clade species *Alitta virens* and the Sedentaria-clade species *Capitella teleta* (de Jong and Seaver, 2016; Kozin et al., 2017)

We previously studied post-regenerative posterior growth and showed that it is highly similar to the normal growth of the worms which occurs during most of its life cycle (Gazave et al., 2013). However, segment addition rate is much higher after amputation than during normal growth (Gazave et al., 2013), indicating that there is a kind of ‘memory’ of the fact that a part of the body has been amputated. Interestingly, we found that segment addition is more rapid if the amputation has been done more anteriorly (Fig. 4D). This could be interpreted in two non-mutually exclusive ways. First, if the amputation is made anteriorly, it implies that more segments are eliminated compared to more posterior amputation planes. The increase of segment addition rate could therefore be related to the fact that the worms are able to ‘sense’ the amount of tissues that have been deleted and to adjust their growth accordingly. Alternatively, there could be positional cues in the worm‘s body and it could be the position of the amputation plane with respect to these cues that defines the rate of segment addition. In agreement with this second possibility, we found that after amputation anterior, parapodia (parapodia are the typical laterally-positioned appendages of annelids) regenerate faster than posterior ones - there is thus a clear anterior to posterior gradient of regeneration speed (unpublished observations). In this case, the same quantity of tissue is removed by the amputation and the only difference is the position along the body axis. What could be the amputation-related signals or positional cues is currently not known. An obvious candidate is the ‘brain hormone’ that promotes growth and inhibits reproduction in nereid annelids such as *P. dumerilii* (Schenk et al., 2016 and references therein). Concentration of this hormone progressively decreases when the worms become older and bigger. Being produced in the head, it is conceivable, but not experimentally demonstrated, that the hormone may have a graded concentration along the body axis, providing an anterior-posterior cue that can be used to control growth. In addition, there are indirect but compelling evidence that posterior amputation may lead to an increased production of the brain hormone (*e.g.*, Clark and Ruston, 1963; Scully, 1964), providing a possible explanation for the accelerated rate of growth after amputation as compared to normal non-post-traumatic growth.

### *Major steps and events during epimorphic and blastema-based* P. dumerilii *posterior regeneration*

Thomas Hunt Morgan, in the early 20^th^ century, proposed the existence of two main modes of regeneration, epimorphosis and morphallaxis, based on whether active cell proliferation is required (epimorphic regeneration) or not (morphallactic regeneration) to ensure proper restoration of the lost body part (Morgan, 1901). Epimorphosis is by far the most common mode of regeneration and often involves the formation of a regeneration blastema (*e.g.*, Sanchez Alvarado and Tsonis, 2006). A blastema is a specialized structure formed upon amputation or injury and which is composed of an outer sheet of epithelial cells that covers an inner mass of mesenchyme-like cells. Regeneration is achieved by the eventual differentiation of cells of the blastema. In annelids, regeneration of the posterior body part is usually epimorphic and involves the formation of a blastema (Özpolat and Bely, 2016). In this article, we demonstrate that it is also the case in *P. dumerilii*. First, using cell cycle genes expressions, EdU incorporations and antibody labelings (Figs. 9 and S7), we show that cell proliferation occurs during most stages of the process and that it increases from stage 2 (2dpa) onwards. Second, we demonstrate the absolute requirement of cell divisions for proper regeneration by showing that worms treated with the anti-proliferative drug hydroxyurea (HU) are unable to regenerate (Fig. 10). Finally, from stage 2 onwards, the posterior regenerated region displays a blastema-like structure with an epithelial superficial layer unsheathing an inner mass of proliferating cells and progressively gives rise in its anterior part to the growth zone and segments, and in its posterior part to the pygidium and its characteristic outgrowths, the anal cirri (Figs. 3, 6 and 9).

Similar characteristic steps and events have been described during epimorphic regeneration in several annelids (Özpolat and Bely, 2016). Hereafter we describe when and how they occur in *P. dumerilii*.

#### Wound healing

as in other annelids, this step is rapidly achieved in *P. dumerilii* (one day or less) and first involves the contraction of muscles at the amputation site, then followed by the formation of a wound epithelium (Figs. 7 and S3). Few proliferating cells are found in the wound epithelium (as well as throughout the worm’s body; Fig. 9), but HU treatments indicate that cell proliferation is not required for wound healing (Fig. 10). Several genes are strongly expressed in wound epithelium (Fig. 6), including genes encoding transcription factors (*Pdum-hox3*, *Pdum-evx*, *Pdum-engrailed*, *Pdum-ap2* and *Pdum-prdm3/16*), signaling molecules (*Pdum-wnt1* and *Pdum sfrp1/2/5*), or RNA-binding proteins (*Pdum-elav*). Several other genes are expressed in mesodermal cells of the segment immediately adjacent to the amputation site (Fig. 6). This includes several ‘stem cells genes’ belonging to the germline multipotency program (GMP), already shown to be expressed during early regeneration stages in other annelids (*e.g.*, Kozin and Kostyuchenko, 2015; Özpolat and Bely, 2015). Interestingly, most of the genes expressed in wound epithelium or adjacent mesodermal tissues, are more strongly or exclusively expressed on the ventral side of the worm, indicating the existence of dorso-ventral cues at this early regeneration stage. *Pdum-elav* shows a particularly striking expression (Fig. 6K), being only expressed in the ventralmost part of the wound epithelium which abuts the ventral nerve cord (VNC), suggesting that its expression might be induced by signals from the VNC. Interestingly, bilateral nerves grows from the VNC from early time points after amputation and underlie the wound epithelium (Figs. 7 and S3), consistent with the hypothesis that *P. dumerilii* regeneration, like that of many other species (reviewed in Boilly et al., 2017a), could be nerve-dependent.

#### Blastema formation

A blastema-like organization is found in *P. dumerilii* from 2dpa onwards. At this stage high cell proliferation is observed (Fig. 9), but HU treatments indicate that this stage can be reached when cell divisions are blocked (Fig. 10). Intense and broad expression of ‘stem cells genes’ are found in the blastema, mainly in its internal mesenchymal-like cells (Fig. 8). An obvious and important question is the fate of the blastemal cells and in particular whether they may have stem cell properties. The expression of homologs of genes such as *piwi*, *vasa*, *nanos* and *myc* may be viewed as suggestive of a stem cell identity for the blastemal cells. However, caution should be taken as these genes are also expressed in non-stem cells, for example during *P. dumerilii* posterior growth in segmental mesodermal and ectodermal progenitors (Gazave et al., 2013). Only cell lineage tracing experiments will allow to safely define the fate of the cells of the blastema.

#### Blastema patterning

As already mentioned, dorso-ventral differences in gene expressions are observed as early as in stage 1 (1dpa). During the following stages, many of the genes we have studied show differential expression along this axis (Figs. 6, 8 and S2). Most stem cell genes are, for example, more strongly and broadly expressed on the ventral side than on the dorsal side. Cell proliferation is also more intense on the ventral side (Fig. 9). While we do not have experimental data in *P. dumerilii* about what could control dorso-ventral organization during regeneration, elegant surgical manipulations in *Nereis pelagica*, which is phylogenetically closely-related to *P. dumerilii*, have provided evidence for a role of the VNC (reviewed in Boilly et al. 2017b). Indeed, when the VNC is absent from the wound site, regeneration occurs, but the aneurogenic regenerated region seems to be only composed of dorsal tissues (*e.g.*, Boilly and Combaz, 1970; Boilly-Marer and Combaz, 1972), suggesting that signals from the VNC are required for the formation of ventral tissues. Anterior-posterior organization of the *P. dumerilii* blastema is also established at early stages. Indeed, at stage 2 (2dpa), some genes, such as *Pdum-wnt1* and *Pdum-dlx* are expressed in the posteriormost part of the blastema while others, such as *Pdu-engrailed* and *Pdum-ap2*, are expressed in its anterior part (Figs. 6 and 8). From stage 3 onwards, a clear demarcation between the posterior pygidium and more anterior growth zone and developing segments is obvious at the levels of gene expression and cell proliferation. How blastema anterior-posterior polarity is controlled is currently not known. An obvious positional cue could be linked to the fact that the anterior part of the blastema is in contact with cells of the last remaining differentiated segments while its posterior part does not.

#### Tissue differentiation

Earliest signs of differentiation, or at least commitment of cells towards a particular differentiated state, are observed at stage 2 (2dpa). Indeed, at this stage, a loosely-organized gut is present (Fig. 2) and some cells in the posterior part of the blastema express *Pdum-twist* (Fig. 6), suggesting that these cells are already engaged to produce pygidial muscles. Phalloidin staining suggests that some of these muscles already start to differentiate (Fig. 7). At stage 3, additional expression of *Pdum-twist* is found in the anterior blastema part, from which segments will be formed, and muscle differentiation occurs at the following stages (Figs. 7 and 8). At stages 2 and 3, cells expressing *Pdum-prdm3/16* are observed and are likely blood vessels progenitors (Fig. 6). Well distinguishable ventral and dorsal blood vessels are observed from stage 4 onwards (Figs 2 and 6). At stage 3, nervous system formation starts as seen by the expression of genes such as *Pdum-pax6* and *Pdum-neurogenin* (Fig. 6). A complex and well organized VNC and nerves are observed both in the developing segment and pygidium from stage 4 onwards (Fig. 7). Finally developing parapodia, expressing genes such as *Pdum-dlx*, are present from stage 3 onwards (Fig. 6).

### *Insights into the cellular origin of the* P. dumerilii *blastema*

A long-standing debate in the annelid regeneration field is the cellular source of the cells of the regenerated region (*e.g.*, Bely, 2014; de Jong and Seaver, 2017; Zattara et al., 2016). This debate covers in fact two distinct questions. One of these questions is whether blastemal cells are produced by divisions of pre-existing stem cells or derive from differentiated cells through dedifferentiation and/or transdifferentiation events, or by a combination of both. The other question is whether the regenerated region is solely produced by cells at, or close to, the amputation site, or may involve the migration of cells from more distant positions. The two aspects are often mixed together by opposing two hypotheses about blastema formation, local dedifferentiation *versus* migration of stem cells from different parts of the body. We however have to keep in mind that migratory cells are not necessarily stem cells and that there could be stem cells located at the amputation site. A third important question is whether different parts of the blastema, ectodermal, mesodermal and endodermal derivatives, have similar or distinct origins, and thus whether pluripotent stem cells or progenitor cells may be involved in annelid regeneration.

One way to distinguish stem cells from differentiated cells is that the former might be proliferating while the latter usually do not. We therefore performed EdU incorporations before amputation to label cells that divide irrespectively of amputation, then amputated the worms and let them regenerate for three or five days (Fig. S9). Only few EdU+ cells are found in the regenerated region at 3dpa and only internal labeled cells are observed. At 5dpa, very strikingly, almost all EdU+ are in the intestinal lining and a large number of gut cells are in fact EdU+. These results indicate that the regenerating intestine mostly or exclusively derives from cells that were proliferative before amputation, whereas mesodermal and ectodermal derivatives do not. A plausible hypothesis is that the *P. dumerilii* gut, like that of other animals, contains stem cells normally involved in the homeostatic replacement of the gut epithelium, but which can, upon injury, also allow its regeneration (*e.g.*, Forsthoefel et al., 2011; Gehart and Clevers, 2015). It is however important to note that our experiments do not demonstrate that ectodermal and mesodermal cells do not derive from stem cells, as these cells could be quiescent before amputation (and therefore not labeled by EdU incorporations) and re-enter cell cycle as a consequence of injury.

To address the question of blastemal cells origin, we designed a second EdU pulse and chase experiment. In this case, we labeled cells whose division was stimulated by amputation by incorporating EdU at 1dpa and we next performed a two-day chase, amputated the worms a second time and let them regenerate for three days (Fig. 11). Importantly, the second amputation was made at three different positions, similar plane than the first one, or one or eight segments more anteriorly. When the same amputation plane was used, meaning that only the regenerated region formed after the first amputation was eliminated and that the differentiated segment that abuts the first amputation site was left in place, the regenerated part formed after the second amputation was full of EdU+ cells, which were found both in the superficial and internal layers of the blastema. In contrast, in the experiments with the two other amputation planes, which have in common that the differentiated segment abutting the first amputation site was eliminated by the second amputation, only few EdU+ cells were found in the blastema after the second amputation and none of these cells were superficial. These results strongly point out a local origin of the blastema, which mostly or exclusively (in the case of the epidermis) derives from cells located in the segment immediately adjacent to the amputation. Our data therefore also argue against the hypothesis of a major contribution to blastema formation of cells migrating from distant segments.

Finally, we tested if primordial germ cell (PGC) progeny could have a role in posterior regeneration. Indeed, these cells were shown to express a collection of stem cells genes, which constitutes the germline multipotency program (GMP), also expressed by the posterior growth zone cells which whom PGCs share a common embryonic progenitor origin (Gazave et al., 2013; Özpolat et al., 2017; Rebscher et al., 2007). PGCs, which produce the germ cells, could therefore also have somatic potentialities in particular in a regenerative context. Using a previously published way to label PGCs (Rebscher et al., 2012) and combining this labeling with amputation, we were able to show that PGCs, at least in our experimental design, do not contribute to blastema formation (Fig. 12). It is conceivable that other stem cells contributing to blastema formation, might exist in the worms’ body, but so far there is no convincing evidence for their existence.

Taken together, our data strongly suggest a mostly or exclusively local origin of the blastema and do not support the hypothesis of a significant contribution of migrating stem cells in posterior regeneration in *P. dumerilii*. Our results also indicate that gut cells have a different origin than mesodermal and ectodermal derivatives. Our data are fully consistent with a set of observations and experiments that were made in the 60s on other Errantia annelids (*e.g.*, Boilly, 1965a,b; Boilly, 1968a,b; Boilly, 1969a,b,c). Careful histological and electron microscopy observations, combined with experimental manipulations, indeed strongly suggested that blastemal cells form through dedifferentiation events and that the different germ layers of the regenerated region have distinct origins. In addition, targeted destructions of cells of different segments have indicated that only the cells of the segment abutting the amputation site were required for regeneration. A very different view has emerged from studies made on Sedentaria annelids, which suggested the involvement of migratory stem cells in regeneration of some species (*e.g.,* Bely, 2014; de Jong and Seaver, 2017; Myohara, 2012; Özpolat and Bely, 2016; Zattara et al., 2016), opening the possibility that profoundly different mechanisms of blastema formation may exist in Errantia and Sedentaria annelids.

### Conclusions

In conclusion, we performed a thorough morphological, cellular and molecular characterization of posterior regeneration in the annelid *P. dumerilii*. We showed that this process involves the formation of regeneration blastema and provide evidence for the importance of cell proliferation for proper regeneration. The cells of the blastema express from early stages a collection of genes known to be markers of pluripotent/ multipotent stem cells, suggesting that the blastema may contain such cells with high potency. We provide compelling evidences that blastema formation is mainly achieved by local dedifferentiation. Taken together, our data pave the way for further analysis of *P. dumerilii* regeneration, and, ultimately, through comparative analyses, for a better understanding of the evolution of regeneration in animals.

## ACKNOWLEDGEMENTS

We thank all Vervoort lab members for helpful discussions and feedback on the manuscript. We are grateful to Aldine Amiel, Bénoni Boilly and Eric Röttinger for helpful suggestions and discussions. We thank Patricia Alvarez-Campos and Thibault Bidolet for help with some *in situ* hybridizations, and Pierre Kerner for providing the *Pdum-Prdm3/16* probe. We acknowledge the ImagoSeine facility (member of the France BioImaging infrastructure supported by the Agence Nationale de la Recherche, ANR-10-INSB-04, “Investments of the future”) and its experienced staff for their assistance in confocal imaging. Animal facility members are thanked for their help with the worm culture. This work was supported by funding from Labex ‘Who Am I’ laboratory of excellence (No. ANR-11-LABX-0071) funded by the French Government through its ‘Investments for the Future’ program operated by the Agence Nationale de la Recherche under grant No. ANR-11-IDEX-0005-01, Centre National de la Recherche Scientifique, and Agence Nationale de la Recherche (grant TELOBLAST no. ANR-16-CE91-0007). JP wishes to thank M. Candás and Xela Cunha-Veira (Estación de Bioloxía Mariña-EBMG, Universidade de Santiago, Spain) and Cati Sueiro and Ada Castro (Servizos de Apóio á Investigación-SAI, Universidade da Coruña, Spain) for their help with use of the microCT and SEM, respectively. Both imaging techniques were financially supported by the project Fauna Ibérica X: Polychaeta VI: Palpata-Canalipalpata I (CGL2014-53332-C5-3-P) from the Ministry of Economy, Industry and Competitiveness (Spain).

